# Allele Surfing Promotes Microbial Adaptation from Standing Variation

**DOI:** 10.1101/049353

**Authors:** Matti Gralka, Fabian Stiewe, Fred Farrell, Wolfram Möebius, Bartek Waclaw, Oskar Hallatschek

## Abstract

The coupling of ecology and evolution during range expansions enables mutations to establish at expanding range margins and reach high frequencies. This phenomenon, called allele surfing, is thought to have caused revolutions in the gene pool of many species, most evidently in microbial communities. It has remained unclear, however, under which conditions allele surfing promotes or hinders adaptation. Here, using microbial experiments and simulations, we show that, starting with standing adaptive variation, range expansions generate a larger increase in mean fitness than spatially uniform population expansions. The adaptation gain results from ‘soft’ selective sweeps emerging from surfing beneficial mutations. The rate of these surfing events is shown to sensitively depend on the strength of genetic drift, which varies among strains and environmental conditions. More generally, allele surfing promotes the rate of adaptation per biomass produced, which could help developing biofilms and other resource-limited populations to cope with environmental challenges.

## Introduction

The dynamics of adaptation has been intensely studied both theoretically and experimentally in situations where the time scales for demographic and adaptive change are vastly separated. Populations can then be treated as either stable or as having an effective population size summarizing the effect of demographic variations on time scales much faster than the adaptive dynamics considered (Muller 1932; Crow and Kimura 1965; Crow and Kimura 1970).

However, demographic equilibrium is frequently disrupted by, for instance, environmental changes, population growth, competition among species and local adaptation (Excoffier et al. 2009). The fate of a genetic variant then both depends on and influences the demography of a dynamically changing population. Consequently, demographic and evolutionary changes can become tightly coupled (Ferriere and Legendre 2012).

Such coupling between ecology and evolution is a particularly salient feature of range expansions (Excoffier and Ray 2008). Many mutations occur in the bulk of a population where they have to compete for resources with their neighboring conspecifics. Mutations that, by chance, arise in a region of growing population densities have a two-fold advantage: They enjoy a growth rate advantage compared to their conspecifics in the slow-growing bulk regions, and their offspring will have a good chance to benefit from future net-growth if parent-offspring locations are correlated. These correlated founder effects, summarized by the term “allele surfing”, lead to complex spatio-temporal patterns of neutral mutations and can rapidly drive mutations to high frequency by chance alone (Edmonds et al. 2004; Klopfstein et al. 2006; Travis et al. 2007; Hallatschek and Nelson 2008).

The importance of allele surfing has been increasingly recognized over the last 10 years (Excoffier et al. 2008; Excoffier et al. 2009; Waters et al. 2013). Allele surfing is believed to be a ubiquitous process in populations that constantly turn over, for instance, by range expansions and contractions, local extinction or expulsion and re-colonization (Hanski 1998; Freckleton and Watkinson 2002; Haag et al. 2005; Taylor and Keller 2007; Arenas et al. 2011). While these features are shared by many populations, they are most evident in microbial communities that frequently expand to colonize new surface regions in the environment or during infections (Cho and Blaser 2012; Costello et al. 2012).

Microbial experiments have shown that in the absence of selection allele surfing creates large mutant clones that are extremely unlikely to arise via neutral evolution of well-mixed populations. Characteristically, these clones take the shape of sectors with boundaries that exhibit characteristic fractal properties (Hallatschek et al. 2007). The random wandering of sector boundaries is a manifestation of genetic drift, as has been demonstrated experimentally in various micro-organisms, including bacteria, single-celled fungi and social slime molds, and under various demographic scenarios (Hallatschek et al. 2007; Korolev et al. 2011; Drescher et al. 2013; Freese et al. 2014; van Gestel et al. 2014).

While allele surfing is well understood in the neutral case, we do not have a comprehensive picture of its adaptive potential. In particular, it is unclear how efficiently pre-existing adaptive variation (Barrett and Schluter 2008) is selected for during range-expansions: Since allele surfing relies on enhanced genetic drift, it reduces the efficacy of selection per generation (Hallatschek and Nelson 2010; Peischl et al. 2013; Peischl and Excoffier 2015). On the other hand, for populations of the same final size, selection has more time to act at the front of a range expansion than in a comparable well-mixed expansion, which could promote adaptation (Hallatschek and Nelson 2010; Zhang et al. 2011; Greulich et al. 2012; Hermsen et al. 2012).

Here, we test whether allele surfing helps or hinders adaptation using microbial competition experiments to measure the efficiency of selection during growth processes. To get a sense of the range of possible evolutionary outcomes, we focus on two extreme cases: spatial range expansions and pure demographic growth of panmictic populations. We find increased adaptation during range expansions and rationalize our quantitative results using theory and simulations.

## Materials and Methods

### Strains and Conditions

Each experiment was performed using a pair of microbial strains that are distinguished by fluorescence and a selectable marker. The fluorescent color difference allows measuring the relative abundance of each strain in competition experiments by fluorescence microscopy as well as flow cytometry. The selectable marker was used to tune the selective difference between the strains in the following way: One strain of the pair, the sensitive strain (called ‘wild type’), grows slower in the presence of a drug, while the other strain, the resistant strain (called ‘mutant’), is largely unaffected. Tuning the concentration of the drug in the medium thus allowed us to adjust the selective difference between both strains. Selective advantages on plates and in liquid culture were measured separately for a range of drug concentrations using the colliding colony assay (Korolev et al. 2012) and flow cytometry (for *S. cerevisiae*), respectively (see Appendix C in Supporting Information), which give consistent results (see supplementary Fig. B1a). Selective differences reported throughout were obtained from linear fits.

*Strains*. We used *S. cerevisiae* strains with W303 backgrounds, where selective advantages were adjusted using cycloheximide. For experiments with *E. coli*, we used both DH5α and MG1655 strains, tuning fitness differences using tetracycline and chloramphenicol, respectively. Additionally, pairs of strains differing only in the fluorescent marker allowed us to perform truly neutral competition experiments (*S. cerevisiae, S. pombe, E. coli*). *S. cerevisiae* and *E. coli* strains with constitutively expressed fluorescent proteins were used to study the dynamics of cells at the front.

A detailed description of all strains and growth conditions is found in Appendix C.

### Main Experiment

Adaptation from standing variation during two types of population expansions (see Fig. 1a): For each pair of mutant and wild type, a mixed starting population of size *N*_i_ was prepared that contained an initial frequency *P*_i_ of mutants having a selective advantage *s*, defined as the relative difference between mutant and wild-type growth rate (Korolev et al. 2012). The population was then grown to final size *N*_*ƒ*_ in two ways, through a range expansion and, for comparison, through uniform growth, and the final mutant frequency *P*_*ƒ*_ was determined. The associated increase in mean fitness 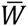 follows as 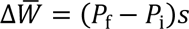.

**Fig. 1:**
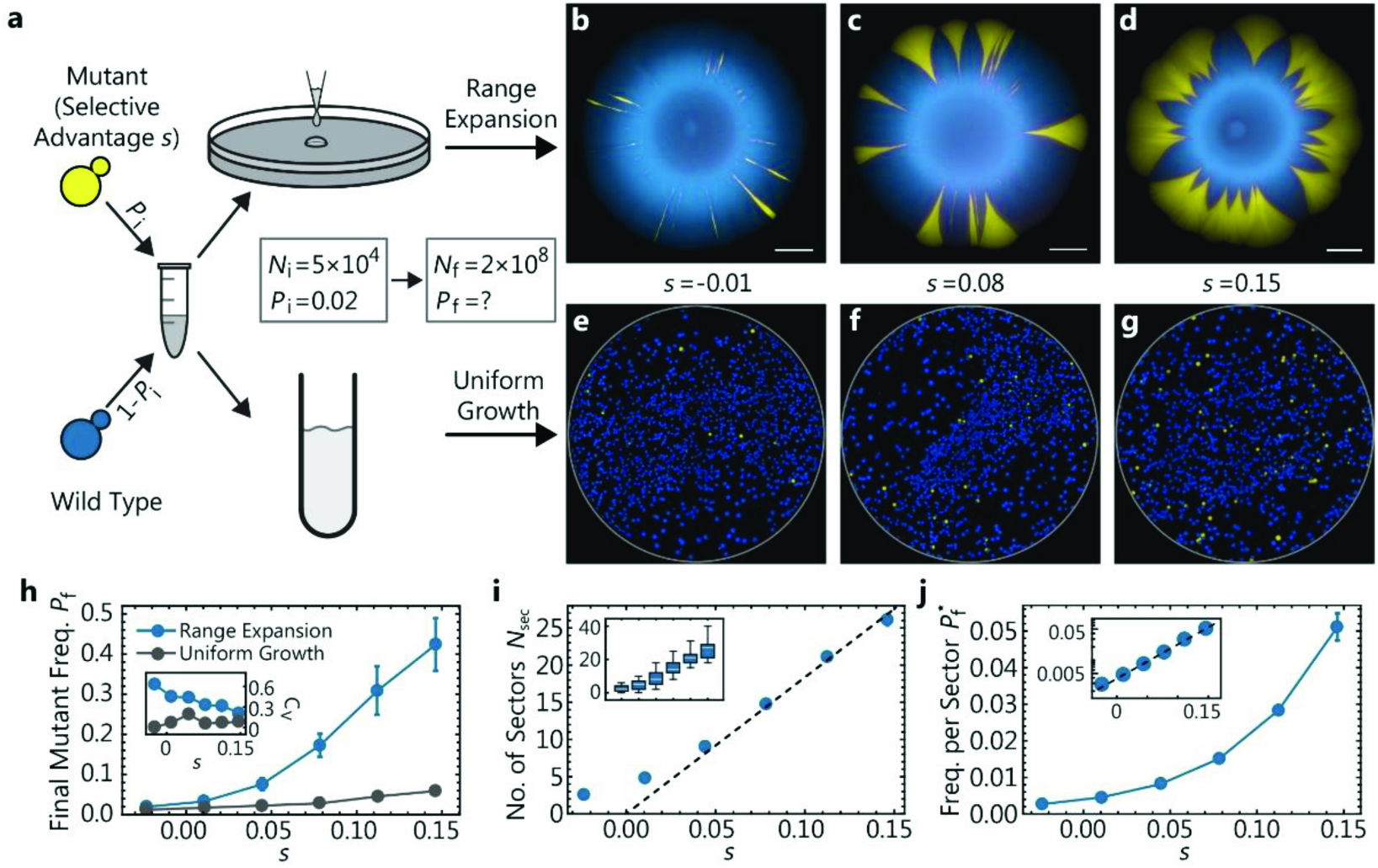
Adaptation from standing variation during a population size increase. Adaptation during the growth of a budding yeast population from an initial size *N*_*i*_ to *N*_*ƒ*_ is studied for two demographic scenarios, Range Expansion and Uniform Growth. (a) Schematic of the experimental assay: Cultures of a wild-type and a faster-growing mutant strain are mixed at an initial mutant frequency *P*_i_ = 0.02. Subsequently, a mixed population of initially *N*_*i*_ = 5 × 10^4^ cells is grown to a final population size of *N*_*ƒ*_ = 2 × 10^8^. The growth process occurred either on agar plates (“Range Expansion”) over the course of 5 days, or overnight under uniform growth conditions (“Uniform Growth”). The selective advantage s of the mutants is controlled by the concentration of cycloheximide, which inhibits the growth of the wild-type cells. The fluorescent microscopy images (b-d) show the distribution of both mutant (yellow) and wild-type (blue) cells at the end of range expansion experiments with selective advantage of s = −0.01,0.08, and 0.15, respectively. Scale bars are 2mm. (e-g) After plating the final populations of the uniform growth experiments, one obtains a distribution of single colonies with a color ratio representing the ratio of mutants to wild type. (h) Final mutant frequency and corresponding coefficient of variation (inset) as a function of selective advantage determined in range expansions (blue, 35 replicates) and under uniform growth (gray, 2 replicates). Notice that the final mutant frequency is larger for range expansions and increasingly so for larger selective differences. (i) Number of sectors *N*_sec_ at the end of range expansions as a function of selective advantage. The inset illustrates the spread of data points as a box plot. (j) Final frequency 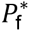 per sector, defined as the area of a single sector normalized by the area of the entire colony, as a function of selective advantage s. The inset displays the same data using a logarithmic axis for the frequency per sector. Only sectors without contact to other sectors were selected for analysis. Error bars are standard error of the mean throughout. The measurements for (h, i, j) were all done on the same 35 replicates per data point.

#### Uniform Growth

Mixtures of cells were grown in well-shaken liquid medium to the desired final population size and the final fraction of mutant cells was determined using flow cytometry.

#### Range Expansion

Colony growth was initiated by placing 2μl of the mixtures onto plates (2% w/v agar) and incubated until the desired final population size was reached. The number *N*_sec_ of sectors was determined by eye; the final fraction *P*_*ƒ*_ was measured using image analysis (see Appendix C for details).

### Cell-Tracking Experiments

To investigate the dynamics of cells at advancing colony fronts, we continually imaged the first few layers of most advanced cells in growing *S. cerevisiae* and *E. coli* colonies between a coverslip and an agar pad for about four hours using a Zeiss LSM700 confocal microscope. The resulting stack of images were segmented and cells were tracked as described in Appendix C.

### Meta-Population Model

To simulate evolutionary change during the different modes of growth, we adapted a classic meta-population model for growing microbial colonies, the Eden model (Eden 1960) (Fig. 2a, Appendix A).

**Fig. 2:**
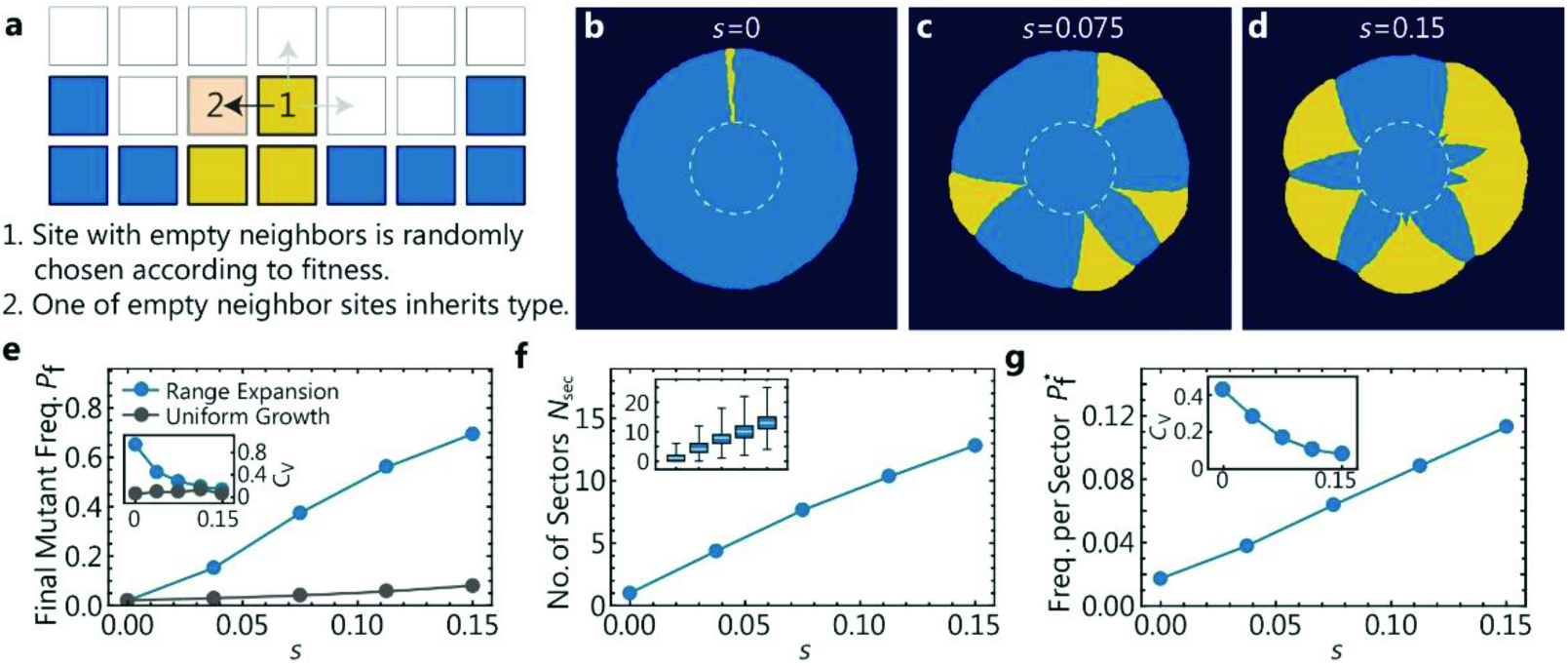
Adaptation from standing variation emerging in a meta-population model of population growth. (a) Illustration of the algorithm underlying our coarse-grained simulations (*Methods*). A lattice site at the population frontier is chosen and copied into an empty neighboring lattice site. The newly occupied site inherits the state of the parent site. (b-d) State of the lattice at the end of three simulations. To mimic our experiments in Fig. 1, we initiated the expanding population as an occupied disk (dashed line) of radius R_i_ ≈ 550 such that a random fraction P_i_ = 0.02 of lattice sites is of the mutant type, and simulated until the final radius R_f_ ≈ 3R_i_ was reached. (e) Final mutant frequency P_f_ and corresponding coefficient of variation C_v_ (inset) as a function of selective advantage s determined in range expansions (blue, 500 simulations per condition) and corresponding simulations of uniform growth (gray, 3 simulations per condition, see *Methods* for algorithm) for the same parameters. Both final frequency and variation are larger for range expansions. (f) Number and standard error of mean of sectors at the end of range expansions as a function of selective advantage for the same simulations. Inset illustrates the spread of data points as a box plot. (g) Frequency per sector 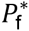, calculated from colonies with only a single sector, which were simulated using a low initial mutant fraction P_i_ = 0.005.

#### Range Expansion

The population spreads on a lattice and each lattice point is in one of three states: empty, wild type or mutant. Growth of the populations occurs by randomly selecting an occupied “source” site with empty neighbors and copying it into a randomly chosen *empty* neighbor site. A mutant is more likely to be picked than a wild-type site by a factor of 1 + *s*. This process is repeated until the colony has reached the final average radius *R*_*ƒ*_ and the final mutant fraction *p*_*ƒ*_ is determined.

#### Uniform Growth

The range expansion simulation was modified such that a target site was an empty site randomly drawn from the entire lattice, rather than from the sites neighboring a given source site.

### Individual-Based Simulations

To study the relevance of microscopic details on the adaptation process, we simulated a growing colony as a two-dimensional collection of sphero-cylinders (rods with hemispherical caps) of various lengths interacting mechanically (see (Farrell et al. 2013) and Appendix A for details). The cells continuously grew (and divided) by consuming nutrients, whose concentration was explicitly computed.

## Results

### The Adaptive Potential of Range Expansions

Our competition experiments in yeast show that when a population grows from a mixture of wild-type cells and faster growing mutant cells by a range expansion (Fig. 1a), it exhibits on average a larger final mutant frequency *p*_*ƒ*_ than a well-mixed population grown to the same final population size *N*_*ƒ*_ 2 ≈ 10^8^ (Fig. 1h). The difference in final mutant frequency between range expansion and uniform growth increases strongly with increasing selective advantage *s* of the mutants. For instance, for *s* = 0.15, mutants make up nearly 50% of the final population (Fig. 1d), in contrast to less than 10% mutant frequency in the well-mixed population. The discrepancy between both growth modes is even more pronounced when we plot the change 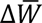=(*P*_*ƒ*_–*P*_i_)*s*. in mean fitness (Fig. B2). Hence, adaptation from pre-existing mutations leads to a much stronger increase in mean fitness in our experiments when a given population increase occurs via the expansion of range margins rather than by a homogeneous density increase.

The spatial distribution of the mutant alleles visible in Fig. 1b-d indicates that the observed adaptation gain of range expansions hinges on the formation and growth of “sectors”. These clonal regions are the footprints of surfing mutants that have locally established at the edge of the range expansion (Hallatschek et al. 2007; Hallatschek and Nelson 2010; Korolev et al. 2012). Sectors contain the vast majority of mutants in the population: If one removes the mutants that reside in sectors from the analysis, or chooses initial frequencies so low that sectors do not occur, the adaptation gain is essentially absent.

Selection has a strong impact on the shape and size of sectors: While a single mutant sector in yeast is stripe-like in the neutral case, it has a trumpet-like shape and can represent a substantial fraction of the total population when the mutants have a selective advantage (compare Figs. 1b-d). The rapid increase of sector size with selective advantage of the mutant strain is quantified in Fig. 1j. For instance, a single mutant sector with selective advantage *s* = 0.15 contains roughly 5% of the total population in our experiments. Under these conditions, a single clonal sector is like an adaptive “jackpot” event that can cause a substantial increase in the mean fitness of the population.

However, the early stages of surfing are a highly stochastic process, and therefore these jackpot events are rare. This is reflected in the rather small number of sectors (proportional to the initial frequency of mutants, see Fig. B3) detected in our experiments. The colonies shown in Fig. 1b-d, for instance, were started with about 10^3^ founder mutants in the inoculum, but only exhibit a handful of sectors (Fig. 1i). The number of sectors varies strongly between replicates (Fig. 1i, inset) and, if the mutants are very infrequent initially, there is a substantial chance that no sectors form (Fig. B4). Importantly, while the number of sectors is generally small, it increases with selective advantage, further contributing to the adaptation gain in range expansions.

### Towards a Minimal Model for Adaptation by Gene Surfing

The population dynamics of our colonies differs from uniform growth in numerous aspects: Cells are delivered to the plate in a droplet, which forms a ring of cells after evaporation (Deegan et al. 1997). The cells start to grow and push each other across the surface of the agar. The population grows at first exponentially, until the growth of the core of the colony slows down due to nutrient depletion behind the front. The further advancement of the front is driven by a layer of proliferating cells (the “growth layer” (Hallatschek et al. 2007; Mitri et al. 2015)) at the edge of the colony (Fig. B5).

While some of these complexities are specific to microbial colonies and biofilms (Nadell et al. 2010), elevated growth rates at range margins combined with local dispersal are the characteristic features of range expansions. To see whether these features alone could reproduce the observed pattern of adaptation, we created a simple meta-population model (Methods), in which the frontier advances by random draws from the demes within the range margins. This simple model has been shown to exhibit universal fractal properties of advancing interfaces (Kardar et al. 1986), which have also been measured in bacterial range expansions (Hallatschek et al. 2007).

As can be seen in Fig. 2, a simulation analog of Fig. 1, the model mirrors our experimental findings: Beneficial mutations have a higher frequency in populations that have undergone a range expansion than uniform expansion. The simulations also reproduce the stochastic formation of sectors and the qualitative dependence of sector number and size on the selective advantage. Thus, the patterns of adaptation seen in our colony experiments seem to originate from the few general features of range expansions that are incorporated in our minimal simulations.

Indeed, we now provide mathematical arguments and individual-based simulations to show how the key features of range expansions conspire to generate the observed adaptation gain; detailed mathematical derivations are provided in Appendix A.

### Qualitative Explanation for Adaptation Gain

We shall begin with a simple, qualitative argument that demonstrates an important difference between range expansions and uniform growth. In a well-mixed population, the mutant frequency grows exponentially with time, *P*_f_ ∝ e^sT^. The number *T* of generations, however, increases only logarithmically with the final population size, *T* ∝ In *N*_f_, such that the mutant frequency changes by *P*_f_/*P*_i_ = (*N*_f_/*N*_*i*_)^s^. In our experiments, this leads to a modest relative change in mutant frequency, e.g., by a factor of 2 for a 6% beneficial mutation over the course of the growth process, which corresponds to about 12 generations. Importantly, the absolute frequency remains well below 1 when the initial frequency is small. Moreover, the final mutant frequency varies relatively little among different replicates, as quantified by the coefficient of variation (Fig. 1h inset). This is because nearly all initially present cells give rise to clones, with similar clone sizes, each corresponding to only a minute fraction of the total population.

In contrast to uniform growth, more generations need to pass to reach the same final population size *N*_f_ in a radially expanding population 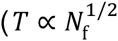 in a radially expanding population, in contrast to *T* ∝ log(*N*_f_) in the well-mixed case). This implies that selection has more time to act during a range expansion, so that one might expect an increased final mutant frequency.

### Adaptation Gain Depends on Sector Shape and Number

The above run-time argument captures the main reason for the adaptation gain, but it ignores two important counterforces: (i) The efficacy of selection is reduced during a range expansion, because the frequency of a selected mutation increases only algebraically with time, in contrast to exponential sweeps in uniformly growing populations. (ii) Only few of the initially present cells give rise to expanding clones. Therefore, to fully understand the adaptive potential of range expansion we must examine the mechanism of sector expansion and formation, the latter being an inherently stochastic process caused by enhanced genetic drift at the front (Hallatschek et al. 2007). Ignoring any interaction between sectors and the small fraction of mutants in non-surfing clones, we can estimate the final frequency P_f_ of mutants by multiplying the number *N*_sec_ of sectors with their relative frequency 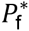 in the population,

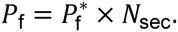

While simple deterministic arguments exist to predict the frequency 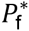 of individual clones, new population genetic theory is required to predict the number *N*_sec_ of sectors. Remarkably, we shall see that the number of sectors is sensitive to microscopic details of the population growth process.

### Final Frequency 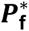 of Expanding Clones

The two boundaries of sectors in radial range expansions are logarithmic spirals (Korolev et al. 2012). These spirals emerge from the origin of the sector at a characteristic opening angle 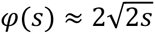 that is set by the selective advantage *s* of the mutant (Hallatschek and Nelson 2010). Up to logarithmic corrections, one therefore expects a final frequency of mutant cells from a single sector to be 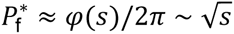 in large colonies (see Eq. (A11) for the full result). This means that a *single* initial mutant can give rise to a macroscopically large clone of order 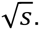 The fractional size of mutant sectors grows even faster in range expansions with straight rather than curved fronts.

### Sector Number *N*_sec_

The establishment of beneficial mutations is generally a result of the competition between random genetic drift and the deterministic force of selection. At the coarse-grained description of clones in terms of sectors, genetic drift manifests itself in the random wandering of sector boundaries, ultimately a result of randomness in the reproduction process (Hallatschek et al. 2007). Balancing the random sector boundary motion with the deterministic sector expansion due to selection, we show in Appendix A (see Eq. (A15)) that the number of sectors is proportional to *s* in two dimensions. Note that although the *s*-dependence of the number of sectors in two-dimensions is identical to Haldane’s classical result “2*s*” for the establishment probability of beneficial mutations (Maruyama 1970; Patwa and Wahl 2008), the proportionality changes in the three-dimensional case to a predicted *s*^3.45^ (Appendix A), which may be relevant to the evolution of solid tumors.

## Modeling the Onset of Surfing

While our minimal model reproduces aspects of the experimental data reasonably well (see Fig. A2), it cannot predict how microscopic details influence the adaptation dynamics. Microscopic details are summarized by a fit parameter, the effective deme size, which enters our expression for the number of sectors *N*_sec_ (Eq. (A19)).

To study *directly* how these microscopic factors influence the number of sectors, we developed an individual-based off-lattice simulation framework for microbial range expansions, where each cell is modeled explicitly as a growing elastic body of variable aspect ratio (see *Methods* and Appendix A). These computer simulations reveal that surfing events result from a complex competition between selection and genetic drift: The probability for an individual cell to form a sector (the surfing probability) increases with selective advantage s but the increase is much faster for colonies with a smooth front line than for colonies with strongly undulating fronts (Fig. 3i). The observed difference between the rough and smooth fronts can be explained intuitively as follows: If a mutant resides in a front region that is lagging behind neighboring wild-type regions, it will likely be overtaken and enclosed by the neighboring wild-type regions, despite its higher growth rate (Fig. 3g). Such “occlusion” events are more likely for rougher fronts, thus increasing the probability that beneficial mutations are lost by chance. In line with this explanation, we find that colonies with rougher fronts also exhibit higher levels of genetic drift, as quantified (Hallatschek et al. 2007) by the lateral (perpendicular to the expansion direction) displacement of lineages from their origin (Fig. 3j). Importantly, we find that front roughness can be strongly influenced by several parameters that can vary among strains and conditions (Fig. A11, Tables A1, A2).

**Fig. 3:**
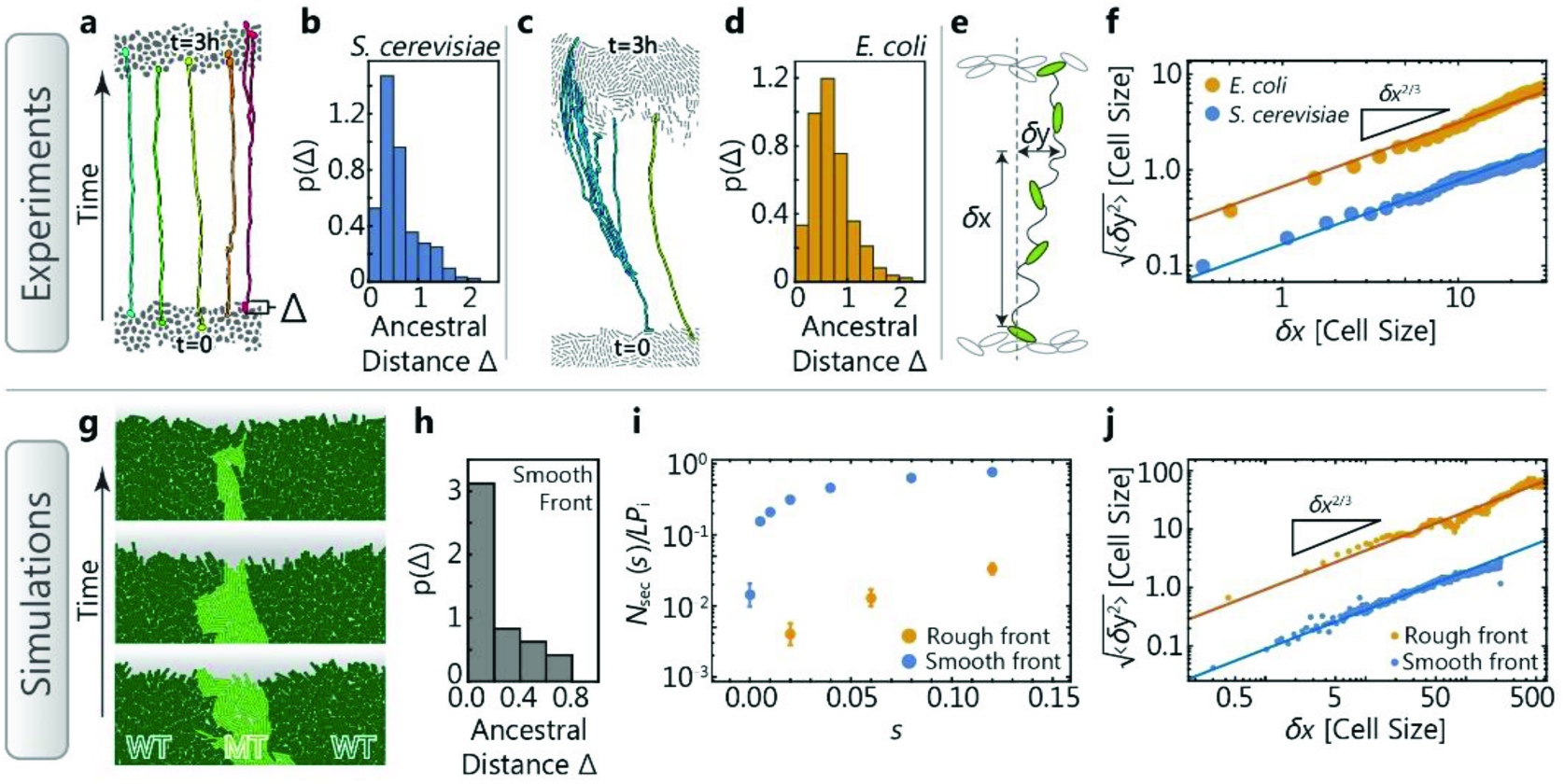
Surfing depends sensitively on location and the strength of genetic drift. Time-lapse microscopy (top row) and individual-based simulations (bottom row) reveal cell-scale dynamics at the front of expanding colonies. (a, c) Segmented micrographs of the initial front (bottom cells) and the front after three hours of growth (top cells) in *S. cerevisiae* (a) and *E. coli* (c) colonies, respectively. Colored lines track lineages backward in time (see also Figs. B8-B10). The histograms in (b, d, h) quantify how surfing success depends on position: The probability density p(Δ) that the lineages tracked for 3 hours back in time lead to an ancestor that had a distance Δ*x* (in unit of cell diameters) to the front. Note the pronounced peak in both experiments (b, c) and simulations (h). (e) Illustration and measurement of the random meandering of tracked lineages. We measure the lateral displacement Δ*y* (in units of cell diameters) a lineage has undergone while moving a distance Δ*x* along the direction of the front propagation, and average 〈Δ*y*^2^〉 over all lineages. (f) Average (root mean square) lateral displacement of lineages in expanding colonies, showing that *E. coli* lineages are fluctuating substantially more strongly than *S. cerevisiae* lineages (absolute value at a given Δ*x*). The lateral displacement in both cases follows a characteristic scaling (slope), as expected for a spatially unbiased growth process with a rough front (Appendix A). These experimental observations can be reproduced in simulations (j) of expanding rough and smooth fronts, respectively. (g) In simulations with rough fronts, surfing beneficial mutations (light green) are frequently occluded by neighboring wild-type domains (dark green). (i) As a consequence, the number of sectors are much lower for rough than smooth fronts, for identical initial mutant frequency *P*_i_ and front length *L*.

Moreover, we find that only mutations that occur very close to the front line have any chance of long-term surfing (Fig. 3h). For our experiments, this implies that only those ancestral mutants have a chance to surf that, by chance, are in the first few cell layers of the dried inoculated droplet. The narrowness of the layer from which surfers are recruited, moreover, makes an important prediction about surfing of *de novo* mutations: Since the width *λ* of the growth layer where mutations occur can be much wider than the average width *d* of the cells in the front line, the *effective* mutation rate μ_eff_ of mutations occurring in the growth layer is the bare mutation rate μ reduced by a factor of *d*/*λ*, which is on the order of a few percent in most microbial colonies. Hence, the vast majority of beneficial mutations are effectively wasted in expanding populations because they occur behind the front line. Therefore, during range expansions with *de novo* mutations, a lot fewer surfing events should be observed than expected for a given mutation rate (as measured by, e.g., fluctuation analysis) and surfing probability (as measured by, e.g., the number of sectors), especially for a thick growth layer. This may contribute to the accumulation of deleterious mutations during range expansions.

## Experimentally Probing the Onset of Surfing

Our individual-based model made two crucial predictions about the early stages of surfing, which we tested in a series of experiments described below.

### (i) Surfing occurs only directly at the front

Control measurements show that the number of surfing events is proportional to the initial frequency (Fig. B3) and not significantly sensitive to the total number of cells, as long as they form a contiguous perimeter around the initial droplet (Fig. B6). These observations are consistent with the hypothesis that surfing events originate in the front region of the colony. To test whether surfers arise in the very first cell layer only, we took time-lapse movies (SI movies 1 and 2) of an advancing front at a resolution that allows us to track lineages backward in time. The resulting genealogies show that only cells at the very front remain as ancestor of future populations. We can extract histograms of ancestor distances from the front (Fig. 3b, d; see also Fig. B10), showing that cells have to be within about one cell diameter to have any chance of giving rise to a successful lineage.

### (ii) The strength of genetic drift influences surfing rates, and is highly variable

We repeated our competition experiments using pairs of *E. coli* (Methods) strains and found up to an order of magnitude differences in surfing probability, i.e., proportion of surfing mutants *N*_sec_/*N*_mut_, for a given selective advantage (Fig. 4). This underscores that the selective advantage of a mutation *alone* has little predictive power over the probability of surfing. The reason is that allele surfing also depends on the strength of genetic drift, which can be estimated from the number of sectors emerging in neutral competition experiments (Fig. 4a, c, e). Fig. 4g shows a clear correlation between the number of surfing beneficial mutations and the number of surfing neutral mutations, for four conditions and different fitness effects. This suggests that measuring the strength of random genetic drift is necessary to predict the efficacy of adaptation.

**Fig. 4:**
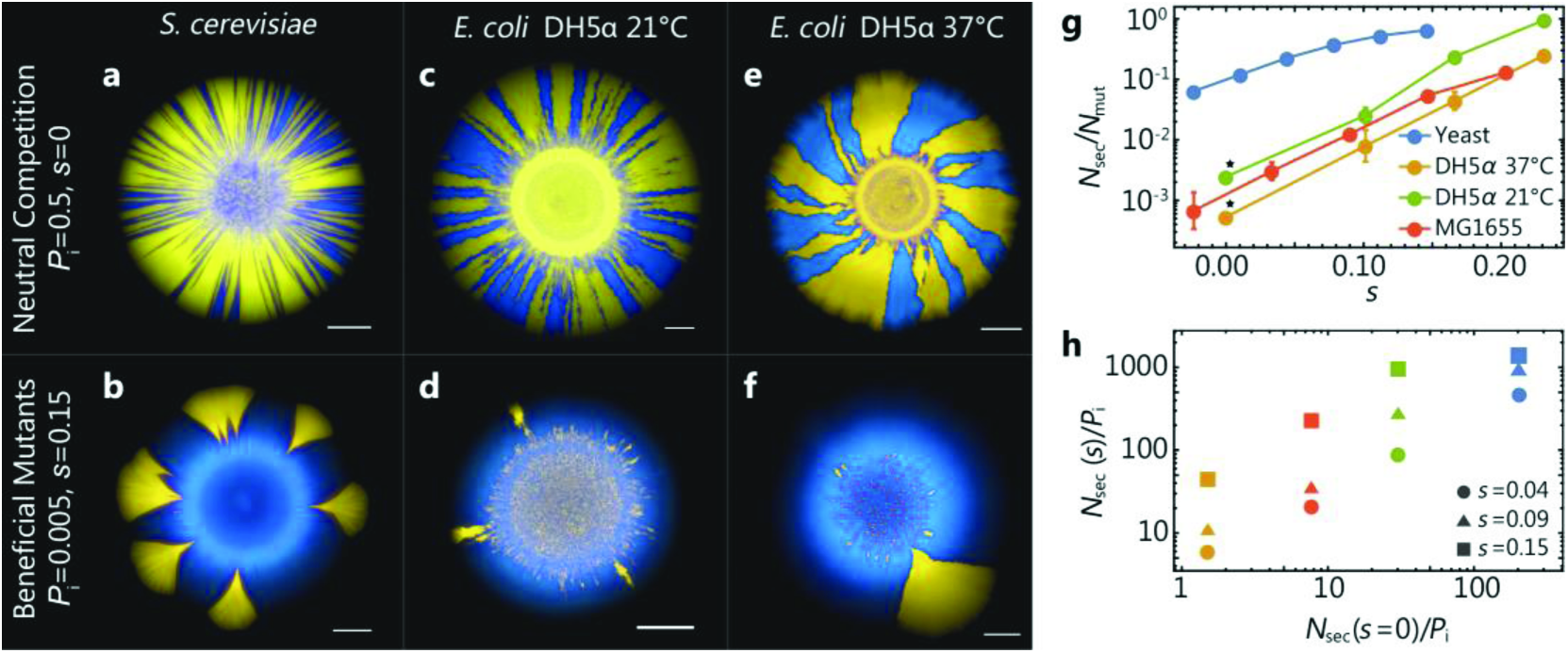
Adaptation during range expansions for different strains and conditions. (a-f) Top row: Images of colonies after neutral range expansions *(Methods)* with an initial mutation frequency of *P*_i_ = 0.5. The number of sectors formed (panel g) and their shape (see Fig. B7) varies between *S. cerevisiae* and *E. coli* and temperature at which colonies are grown. The bottom row shows corresponding range expansions when mutants have a selective advantage of s ≈ 0.15, at low initial mutation fraction of *P*_i_ = 0.005. Scale bars are 2mm in each image. (g) The number *N*_sec_ of sectors normalized by the number *N*_mut_ of mutant cells in the outside rim of the inoculum as a function of the selective advantage of the mutants for different species, strains, and growth conditions (about 35 replicates per data point). The asterisk (*) denotes the use of the neutral strain pairing as opposed to the mutant-wild-type pair. (h) The number of sectors *N*_sec_ normalized by the initial fraction *P*_i_ against the normalized number of sectors in the neutral case shows a clear correlation between neutral dynamics and the surfing probability of advantageous mutant clones: weaker genetic drift (more sectors in neutral competitions) is indicative of a higher surfing probability. Panel (h) is obtained by interpolating data from panel (g) for the selected values of *s*.

The difference between strains can partly be understood from time-lapse movies of the colony growth at single-cell resolution (SI movies 1 and 2). While cell motion perpendicular to the front direction is limited in yeast colonies, there is strong dynamics within the *E. coli* front. Tracking the cells through 3 hours of growth elucidates the difference in cellular dynamics, as shown in Fig. 3a and c. We quantify this observation by measuring the cells’ lateral displacement (Fig. 3e-f, Appendix C), which is about an order of magnitude stronger in *E. coli* compared to budding yeast, explaining (at least part) of the difference in genetic drift. The same effect can be observed in computer simulations of the individual-based model (Fig. 3i, j).

While it may not seem surprising that genetic drift varies somewhat (though not an order of magnitude) between taxa due to differences in the reproductive process, we also found that the level of genetic drift varies among different growth conditions for the same species. Fig. 4c-f show the results of competition experiments between two differently labeled but otherwise identical *E. coli* strains (DH5α background) at two different incubation temperatures. Notice that the neutral sectoring pattern undergoes a striking change: While only few sectors can be observed at 37°C, many spoke-like sectors arise at 21°C. Importantly, surfing probabilities varied, as predicted, with observed variations in the strength of genetic drift: repeating the establishment experiments at lower temperatures shows that the number of established clones indeed increased for smaller amounts of genetic drift (Fig. 4g, h).

## Discussion

Laboratory evolution experiments usually investigate the rate of adaptation *per unit time*. This is the relevant quantity when resources are abundant or replenish faster than they are consumed, as for example in a chemostat (Kawecki et al. 2012).

By contrast, in our experiments we have compared the adaptive outcome of two types of population expansions, range expansion and uniform growth, under the condition that both types lead to the same final population size, no matter how long it may take. Thus, we have effectively measured the rate of adaptation *per cell division* or, equivalently, *per biomass produced*. We believe this is the crucial comparison when population growth is resource-limited, which may arguably apply not only to microbial biofilms (Stewart and Franklin 2008; Mitri et al. 2015), but also to various other types of natural populations, including tumors, and spreading pathogens (Lee 2002; Ling et al. 2015).

Our experiments show that, starting from standing adaptive variation, range expansions generate a larger, often much larger, mean fitness increase in microbial communities than equivalent uniform population expansions. In essence, this results from the effective serial dilution of the pioneer population, generated by the fact that the offspring of pioneers tend to be the pioneers of the next generation. As a consequence of these spatio-temporal correlations, selection can act over more generations at the front of a range expansion than in a uniform expansion.

However, because the relevant pioneer population is small, sampling effects (genetic drift) are important: The gain in adaptation comes in partial sweeps, visible in our experiments as large “sectors”, which represent successfully surfing alleles. The total adaptation gain during a range expansion depends on both the number of sectors and the size of sectors. While the shape of sectors simply reflects the selective advantage of the mutants, the stochastic number of sectors is a result of the competition between selection and (strong) genetic drift in the pioneer population.

Thus, predicting the number of sectors, and ultimately the rate of adaptation in population expansions, requires a measurement of both the strength of selection and genetic drift. In microbial experiments, the strength of genetic drift, which is related to the front roughness, can be measured by neutral mixing experiments with fluorescently labeled strains. Such measurements show that the strength of genetic drift varies by orders of magnitude among strains and conditions like growth medium or temperature, affecting surface roughness, growth layer width, or cell shape, as illustrated in Fig. 5. Thus, changes in the microbial growth processes can strongly influence the adaptive potential of range expansions via their impact on the strength of genetic drift. This may be important, for instance, for adaptation in developing biofilms with their complex surface properties (Xavier and Foster 2007; Drescher et al. 2013), and could be tested in flow chamber experiments.

**Fig. 5:**
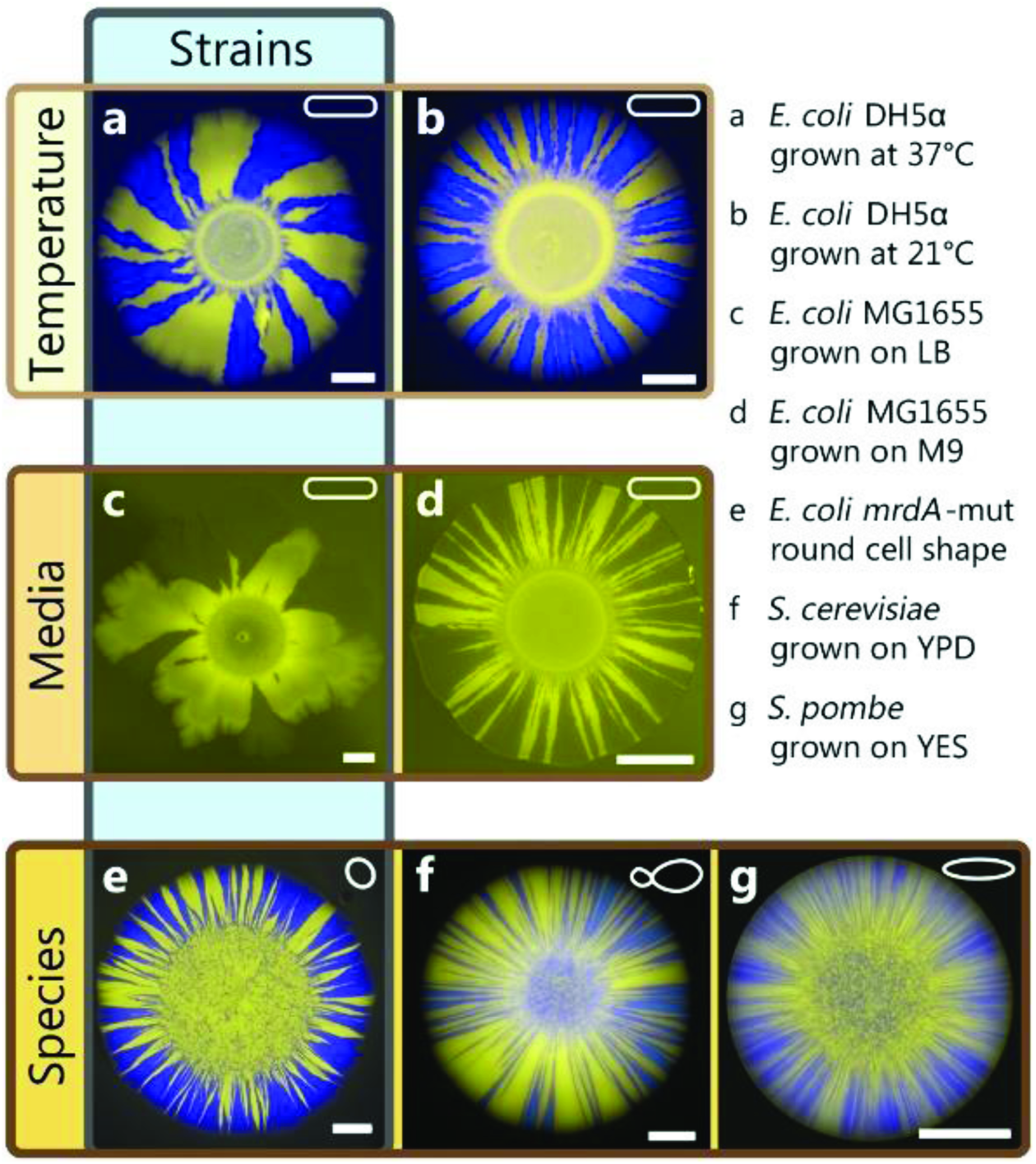
Variability of genetic drift across species, strains, and environmental conditions. Each image shows a colony of two neutral strains grown with a starting frequency *P*_i_= 0.5. Colored frames indicate the main differences between images. *E. coli* colonies (a-e) exhibit fewer sectors and are less regular than yeast colonies (f-g), which produce many sectors. Environmental factors, in particular temperature (a-b) or composition of media (c-d) also influence the strength of genetic drift. Even for identical conditions, different *E. coli* strains exhibit varying morphologies and sector numbers: For example, mutations influencing cell shape (e) may leads to straighter sectors boundaries and more sectors, although cell shape alone does not accurately predict the strength of genetic drift (compare *E. coli* (a-d) and *S. pombe* (g), which are both rod-shaped). All scale bars are 2mm.

Our results underscore the adaptive potential of allele surfing: Although, as was found previously in the neutral case, allele surfing is a rare event that depends on enhanced genetic drift at the frontier (Hallatschek et al. 2007), it becomes more likely as the selective advantage of the mutation increases. Nevertheless, out of the preexisting mutant population only few mutants manage to establish and surf at the frontier. The ones that do, however, leave a strong mark on the population as a whole; driven by selection, their descendants sweep to high frequencies in the population.

In other words, allele surfing turns a population expansion into a high-paying evolutionary slot machine (Luria and Delbrück 1943): The expected gain in fitness is high on average but it relies on rare surfing events controlled by the competition of genetic drift and selection. Range expansions can thus lead to large evolutionary change if these jackpots events do occur. By contrast, well-mixed populations lead to a homogeneous growth of all cells, resulting in less overall change in frequencies. As our experiments have focused on standing genetic variation, they have ignored the impact of spontaneous mutations occurring during the population expansion. Enhanced genetic drift at expanding frontiers is expected to promote the genetic load due to new deleterious mutations (Travis et al. 2007; Hallatschek and Nelson 2010; Peischl et al. 2013; Lavrentovich et al. 2015), which may lead in extreme cases to a slowdown of the population expansion, for instance when “mutator” strains are involved. Thus, enjoying an adaptation increase from a range expansion may require a sufficiently low rate of deleterious mutations.

Strikingly, our expanding colonies shifted from a predominantly wild-type to a largely resistant population under quite weak selective pressures. We hypothesize that adaptation by allele surfing could be a general mechanism for efficiently shifting the balance between pre-existing types after an environmental change. Moreover, a proposed connection (Lambert et al. 2011) between drug resistance in bacterial communities and malignant tissues suggests that similar effects could be at play in solid tumors that harbor standing variation prior to drug treatment.

Allele surfing may also help explain the efficient adaptation seen in some cases of species invasions, such as in cane toads, which developed longer legs in the course of the invasion of Australia (Phillips et al. 2006). Although we do expect our results to carry over to more complex scenarios, sex, death, recombination, dominance, and heterogeneities in resources and selection pressures may significantly complicate the dynamics. Key differences could arise, for instance, if mutants do not have an expansion velocity advantage, but are instead merely outcompeting the wild-type individuals within already occupied regions. In this case, we expect sectors to reach substantially lower frequencies than in our experiments.

Adaptation by gene surfing matches the pattern of a “soft” selective sweep (Hermisson and Pennings 2005; Barrett and Schluter 2008), in which multiple adaptive alleles sweep through the population at the same time, however with a unique spatial structure. Although these sweeps can be strong, as seen in our experiments, they may be hard to identify in population genomic studies when they carry along different genomic backgrounds. However, as sequencing costs drop further and spatial sampling resolution increases, the genomic signal of these localized soft sweeps may become directly discernable.

## Acknowledgments

We thank Melanie Müller and the lab of Andrew W. Murray for providing us with the unpublished strain yMM9. We thank Laurent Excoffier and Stephan Peischl for helpful discussions and a critical reading of the manuscript. Research reported in this publication was supported by the National Institute of General Medical Sciences of the National Institutes of Health under Award Number R01GM115851, and by a Simons Investigator award from the Simons Foundation (O.H.). The content is solely the responsibility of the authors and does not necessarily represent the official views of the National Institutes of Health. This research used resources of the National Energy Research Scientific Computing Center, a DOE Office of Science User Facility supported by the Office of Science of the U.S. Department of Energy under Contract No. DE-AC02-05CH11231.

## Appendix A: Theory and Simulations

**1 Coarse-grained simulations and analytical results**

**1.1 Simulation algorithm**

We simulate range expansions using a metapopulation model on a lattice, similar to the Eden model. Initially, the central site of an empty lattice is filled with a single cell. In each time step, a cell with at least one empty neighboring lattice site is randomly chosen to divide into one of the empty sites in its 4-site neighborhood. If there are mutants in the colony with a selective advantage s, the algorithm first randomly chooses whether to forward the wildtype or mutant population, where the mutants are chosen with probability

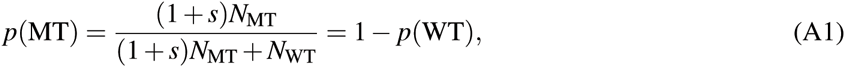
where N_MT_ and N_WT_ are the number of mutant and wild type site having empty neighbors.

**Standing variation**. The colony is first grown to a radius *R*_*i*_ (by running the simulation 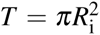 steps; for Fig. 2, *T* = 10^6^) of only wild types. Then, filled lattice sites are randomly populated with wild types and mutants at a specified ratio *P*_i_. The colony is grown a total of 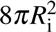 time steps, i.e., to a final radius of about 2.8 x *R*_*i*_. This corresponds roughly to the radial increase in our experiments.

**Figure A1:**
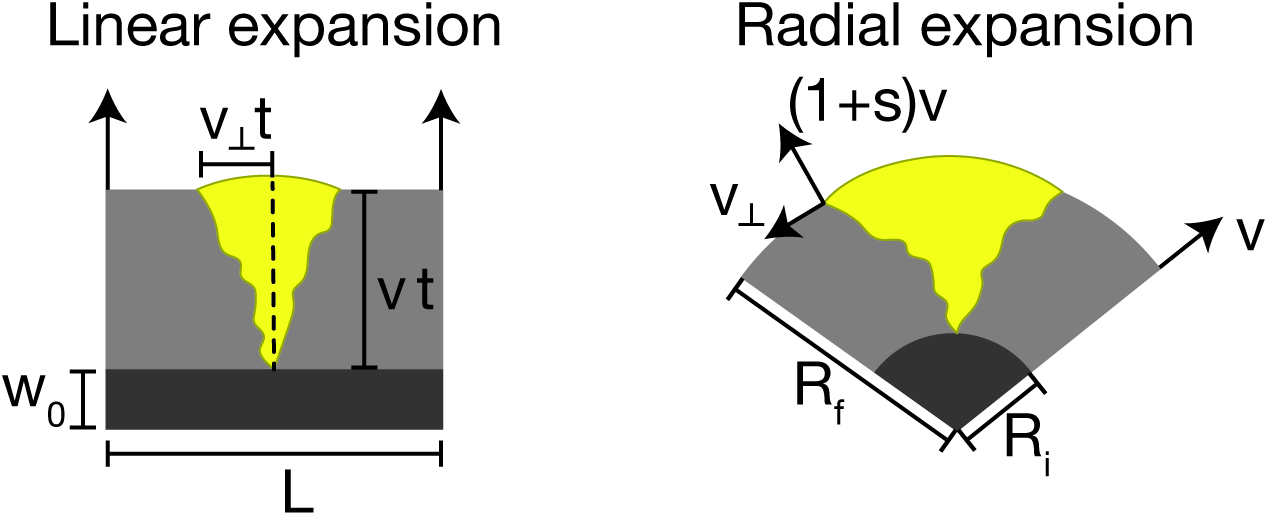
Sketch of the expansion of a sector in a linear (left) and a radial range expansion (right). While sectors have a constant opening angle φ in a linear expansion, their boundaries form logarithmic spirals in the radial expansion case, enclosing an angle φ that increases logarithmically with the radius *R*_*ƒ*_ (cf. Eq. (A9)).

For the scaling function below, Pi was varied between 0.02 and 0.005 to minimize interaction between sectors. Sectors were counted by identifying all mutant clones that have at least one member with at least one empty neighboring lattice site at the end of the simulation.

**De novo mutations**. Instead of starting from a mixture of wild type and mutant sites, we can allow for spontaneous mutations. Populations are grown from a single individual, and every new individual has a chance *JJL* of converting to the mutant type, having an advantage *s*. Here, we do not consider back mutations.

**Long range jumps**. To interpolate between the well-mixed and the colony case, we simulate long range jumps by following Ref. [1]. A random number *Y* between 0 and 1 is drawn and transformed to a jump length *r* by computing

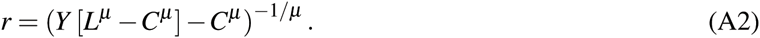

Here, *L* and *C* specify the maximum and minimum jump length. The new variable *r* is distributed as a truncated power-law with a power-law tail, i.e., *p*(*r*)∼r^−μ^. To allow for long range jumps, we employ periodic boundary conditions. In addition, an angle φ is drawn between 0 and 2*π*. In every step, a random lattice site *(xi,yi)* is chosen and the jump attempted to the lattice site located closest to (*x*_*i*_ + *r*cos φ, *y*_*i*_ + *r* sin φ); if the site is empty, it is filled, otherwise a new site is chosen. Only successful jumps forward the time variable, such that exactly one jump happens in each time step. After *T*_*i*_ steps, mutants are introduced by randomly mutating each filled lattice with a probability equal to the desired ratio of wild type to mutant cells. Thus, the initial frequency of mutants is stochastic, mimicking the situation in real experiments.

**1.2 Final mutant frequency**

In the following, we refine the scaling arguments given in the main text to explain the increased adaptation gain in range expansions. To reach the same final population size, a larger number of generations at the front of a range expansion is necessary, allowing selection to act for longer, compared to exponentially growing populations. Yet, selection is weaker at the advancing front in the sense that a selective advantage *s* does not lead to an exponential increase in frequency like it does in well-mixed populations. Nevertheless, we argue below that the former effect is in general stronger than the latter, leading to a net increase in adaptation gain.

**Well-mixed population**

Starting from *N*_*i*_ initial cells, of which a fraction *P*_i_ are mutants, the number of mutant cells *M*(*t*) at time *t* (in generations) is *M*(*t*) = *P*_i_*N*_*i*_2^(1+*s*)*t*^. To reach final population size *N*_f_, it takes *t* = log_2_(*N*_f_/*N*_*i*_) generations, hence, *M*(*t*) = *P*_i_*N*_*i*_(*N*_f_/*N*_*i*_)^1+s^. The final mutant frequency thus becomes

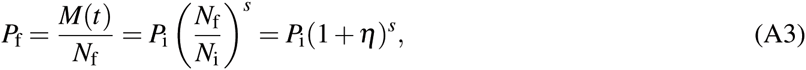
where we have defined the fold change η = *N*_f_/*N*_*i*_ - 1 of the total population size. The adaptation gain in a well-mixed population can be quantified through the fold change R_WM_ of the mutant frequency

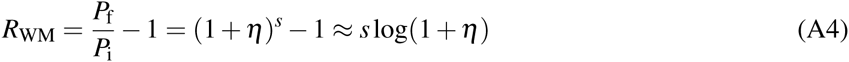
for *s* ≪ 1. For small η, this reduces to R_WM_ ≈ η*s*.

**Flat front range expansion**

Start from a region of (constant) height *L* and width *w*_*0*_, containing *N*_*i*_ = *Lw*_*0*_ individuals (see sketch in Fig. A1, left). We assume that the width grows at speed v, and sector size increases with perpendicular velocity *v*⊥ = 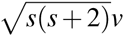 [5]. The final mutant population size is composed of the size of (roughly triangular) beneficial sectors times their number, plus the neutral contribution, i.e.,

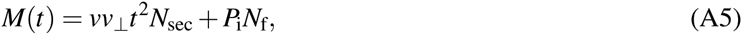
where we have ignored fluctuations of the sector boundaries as well as the typically small number of mutants in non-surfing clones. The number of generations to reach final size *N*_f_ is *t* = (*N*_f_–*N*_*i*_)/*vL* = η*N*_*i*_/*vL*. Plugging this into *M*(*t*) and dividing by *N*_f_ to find the final mutant frequency, we get

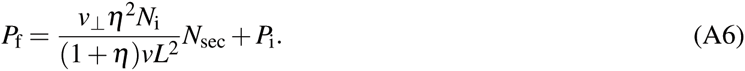

The number of sectors can be estimated as *N*_sec_ = *LP*_*i*_*u*(*s*), where *u*(*s*) is the (unknown) probability to form a sector per individual at the front, i.e., the surfing probability. Hence, we obtain the fold change *R*_*FF*_ = *P*_*ƒ*_/*P*_i_ −1 in the flat front case as

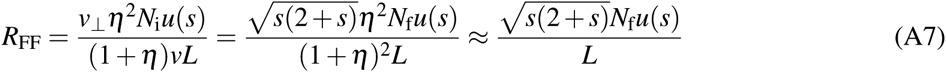

for η ≫ 1. Thus, for a final population size much larger than the initial population size (as is the case in our experiments), the size of the adaptation gain *R*_*FF*_ depends critically on the surfing probability *u*(*s*). This indicates that a purely deterministic treatment is not appropriate to understand adaptation during range expansions. Adaptation crucially hinges on sector formation. Nevertheless, for some fixed *s*, Eq. (A7) shows that in the long run, range expansions will always produce a larger adaptive outcome than exponentially growing populations as the linear scaling of *R*_*FF*_ with *N*_f_ will eventually overtake the logarithmic scaling of *R*_*WM*_.

**Radial expansions**

The situation is less straightforward in a radial expansion, as the shape of sectors is influenced by both inflation and selection. Their shape and size can be understood from simple geometrical arguments [5, 6], which we replicate and extend here.

Mutants grow faster into the expanding territory by a factor of 1 *+s* (see sketch in Fig. A1, right). This speed difference together with the requirement of continuity of the colonial edge enforces a fixed speed at which mutants expand (wild-types retract) along the colony edge. The transverse expansion speed 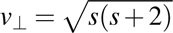 (in units of the wild-type front speed) follows from equating the speed of radial growth in both compartments (1 vs. 1 + *s*). As a consequence of the transverse expansion of the two sector boundaries, the opening angle φ of the sector increases with radial distance according to

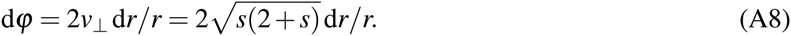

Integration yields a logarithmic increase with radius,

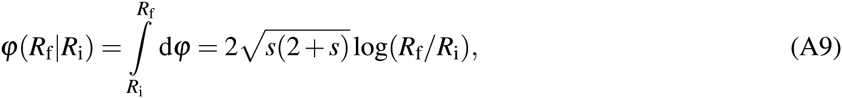
as was already shown in Ref. [6]. Assuming large sectors such that the initial period of sector formation is negligible, the final frequency of the sector is obtained by integration,

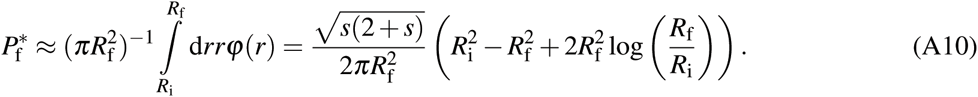

Defining the fold change in the population size η through 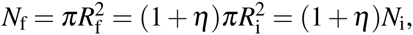 we get

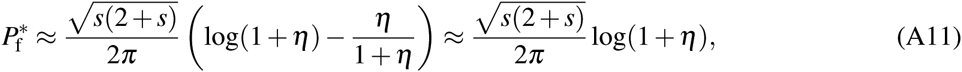
where we have assumed η ≫ 1 in the final step. We again define *N*_sec_ ≡ *LP*_*i*_*u*(*s*), where here *L* = 2π*R*_*i*_, and obtain the fold change in mutant frequency for radial expansions as

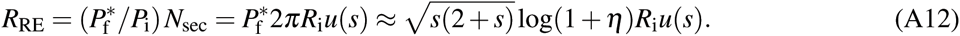

Comparing this to the well-mixed result we obtain

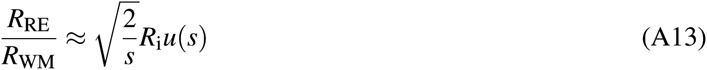
for 0 < s ≪ 1. As in the flat front case, the surfing probability enters in determining the adaptation gain increase of the range expansion compared to well-mixed population. The crucial difference to the flat front case lies in the fact that *R*_RE_/*R*_WM_ is independent of *N*_f_. It is thus ultimately the number of sectors that elevates the adaptation gain in the radial range expansion over the well-mixed one. Therefore, a detailed understanding of the establishment of sectors is necessary. Previous calculations of the surfing probability in boundary-limited radial range expansions have predicted 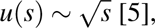 [5], which would remove the dependence of *R*_RE_/*R*_WM_ on *s*. As we have seen in Fig. 1I, this is not the case in our experiments, where we find instead *u*(*s*) ∼ *s*. This linear dependence is reminiscent of the classical Haldane result, but we show below that this similarity is fortuitous and can in reality be traced back to surface growth properties of colonies.

**Figure A2:**
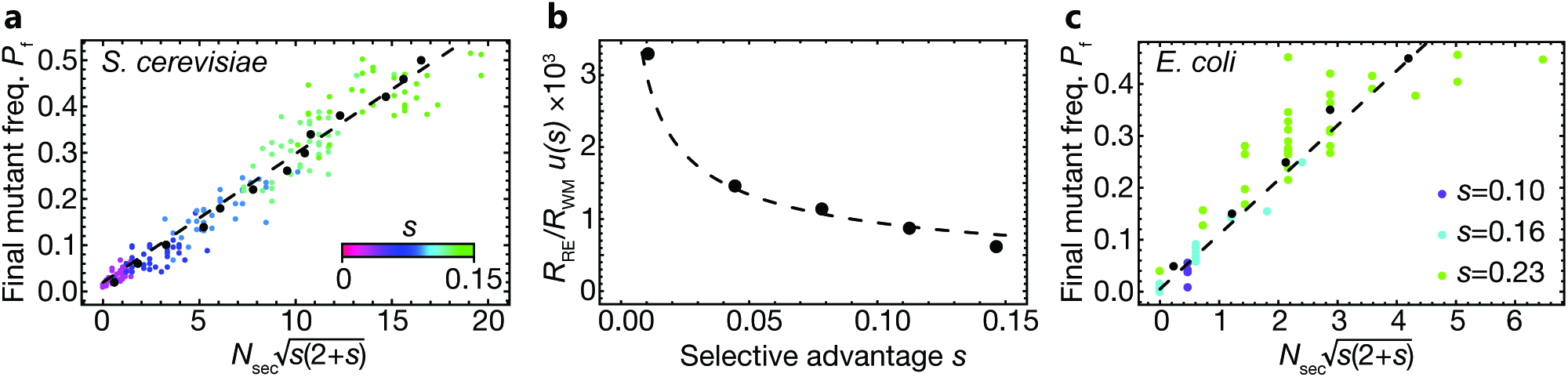
Validating the minimal model with experimental results. (a) Final mutant frequency *Pf* in *S. cere-visiae* colonies, as a function of 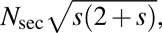 which exhibits the predicted linear scaling (see eq. (A12), dashed line). Each dot corresponds to a colony with mutant selective advantage given by the color legend. Black dots are average values over mutant frequency bins of width 0.04. (b) The ratio between *R*_RE_ and *R*_WM_, normalized by the surfing probability of a single clone, as a function of *s* is consistent with the predicted 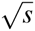 scaling (Eq. (A13), dashed line). (c) Final mutant frequency *P*_*ƒ*_ in *E. coli* DH5α colonies, as a function of 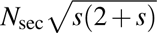 Black dots are average values over mutant frequency bins of width 0.1.

**Validating the minimal model**

Our result thus far neglects the fact that the mutant sectors have a larger area than a wild-type sector of the same opening angle because it bulges outward at the colony rim. Numerical estimates of the correction show that this contribution is not always negligible, especially for large *s*. To improve the calculation, one could account for the fractional area of the circular cap associated with a mutant sector of given opening angle and selective effect. In addition, the sector shape computed above is only valid far from the inoculum, where initial stochastic effects of sector formation no longer impact the shape of the sector. Lastly, in some of our experiments, sectors collide and hence cover a slightly smaller area than if they had grown undisturbed.

Nevertheless, we can compare our experimental data to the theoretical prediction. Fig. A2 (left) shows the final mutant frequency 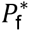 as a function of the number of sectors, for each colony, multiplied by the 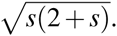 The averaged data (black dots), fall on a line, as predicted by Eq. (A12).

In addition, our results predict that the ratio *R*_RE_/*R*_WM_ of the adaptation gain from a range expansion and uniform growth should scale as 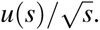 Normalizing by the experimentally measured surfing probability *u*(*s*) ≈ *N*_sec_/2π*R*_i_, we recover the predicted scaling 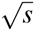 see Eq. (A13) and Fig. A2 (right).

**1.3 Number of surfing clones**

The deterministic calculations for the adaptation gain in range expansions hinge on the likelihood of the formation of sectors. Computing the number of sectors, or “surfing clones”, is a stochastic problem that involves the fluctuation statistics of growing microbial colonies. While these fluctuations are complicated to derive microscopically, their overall scaling behavior is well understood, allowing us to derive the relationship between the number of sectors, the selective advantage and the initial conditions of the population.

**Linear fronts, standing variation**

Consider first the case of a linear front with a small initial fraction *P*_i_ ≪ 1 of mutant sites. As the population edge advances, the extinction and growth of a mutant sector will be dominated by genetic drift as long as the lateral size *l*⊥ of the sector is smaller than some characteristic size 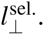 Once a sector has reached this size 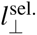, selection takes over and it is unlikely that the sector goes extinct (at the front). Thus, we may call 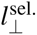 the establishment size for surfing. If we knew 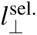 we could estimate the surfing probability by a martingale argument, as follows. Since the dynamics of a sector below size 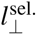 is neutral, all of the 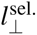 front ancestors have the same chance to generate a clone that drifts up to size 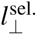 or larger. Thus, we can estimate the probability *u*(*s*) of a mutant clone to surf as

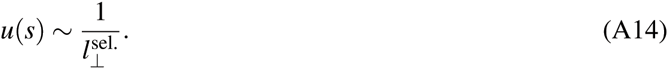

**Figure A3:**
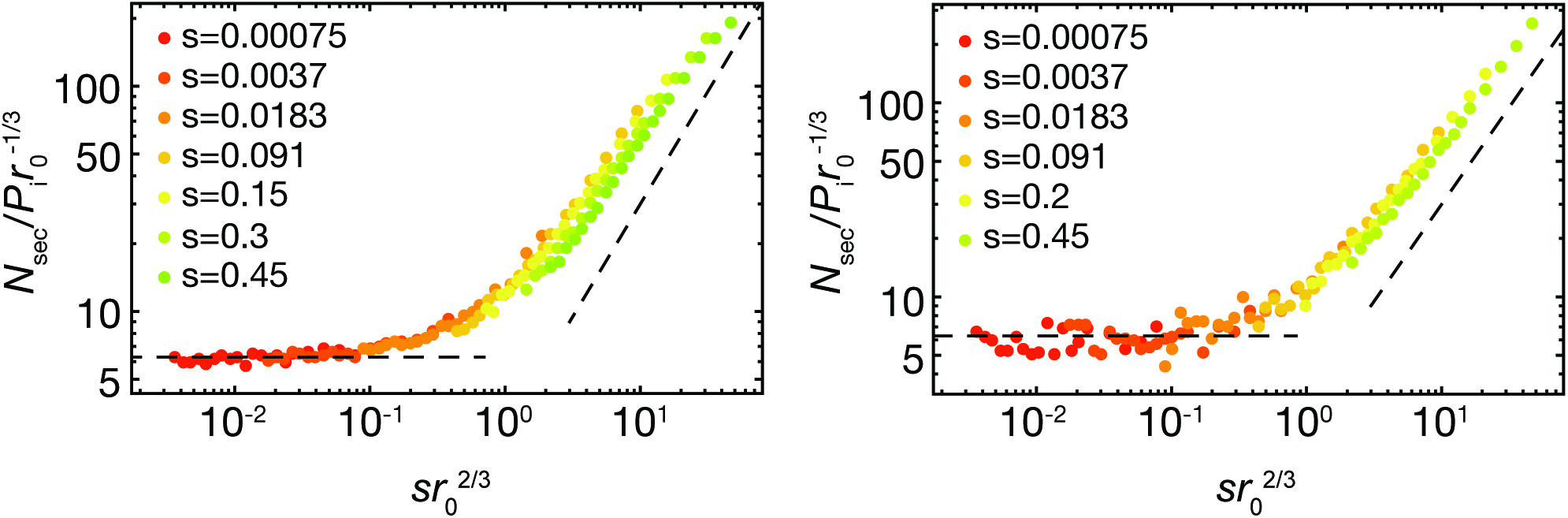
Number of sectors in Eden simulations with standing variation follows a scaling form. Left: Scaling function starting from a droplet of radius *r*_0_. Here, we chose the initial mutant frequency as *P*_i_ = 0.02 like in the experiments. Lines are guide to the eye, showing the predicted constant and linear regimes. At large s, deviations from a linear scaling become visible, because sectors inevitably begin to interact. Right: Scaling function for *P*_i_ = 0.005, this time grown from a single cell to a population of (average) radius r_0_, then inserting mutants at ratio *P*_i_, for a wide range of *r*_0_. The scaling function is virtually indistinguishable from that for flat initial conditions. The plot legend explains the color code for the selective differences.

Since we begin with a fraction of *P*_i_ initially mutated sites, we expect a number 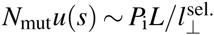 of successful surfing events, where *L* is the length of the front. Note that one has a simple linear dependence on *P*_i_ only for small 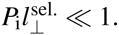 For larger *P*_i_, sectors may overlap when they are still smaller then their establishment length, leading to (predictable) deviations from the observed scaling: The actual number of surfing events will be smaller than estimated.

The establishment length 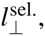, and consequently the number of surfers, is controlled by a competition between selection and genetic drift. The smaller *s*, the larger the sector needs to become, by chance, for selection to take over genetic drift. Genetic drift in our colonies depends on the roughness properties of the colony edge: The rougher the front, the larger the stochastic evolutionary outcomes are. To estimate the establishment length 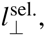 we need to invoke the universal fractal properties of Eden fronts which are in the Kardar-Parisi-Zhang (KPZ) universality class [7]. Conditional on survival, a neutral sector reaches size *l*⊥, roughly, after a time of order 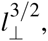, a KPZ prediction that was confirmed in Ref. [8]. Thus, the magnitude of the speed of growth of the width of a sector due to random genetic drift scales as 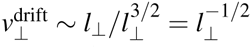 (again in units of the wild-type front speed). Selection on the other hand increases a sector width linearly in time according to a constant speed 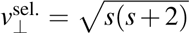 [5]. Both speeds balance at a length scale of 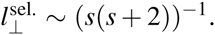 Genetic drift dominates 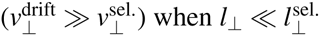 and selection dominates 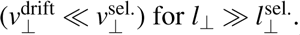 Knowing the establishment length now allows us to predict the scaling of the number of sectors 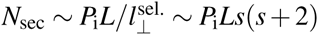.

**Figure A4:**
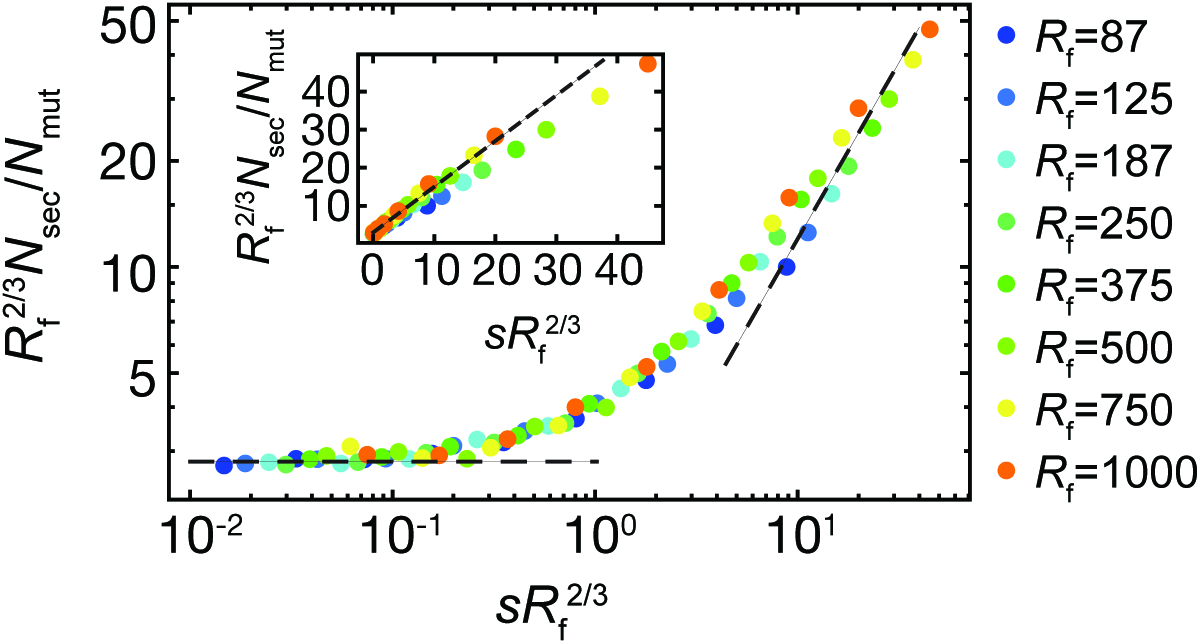
Number of sectors for colonies with de novo mutations obeys a scaling relation, for a wide range of selective advantages s and final radius *R*_f_ (see color legend). The number of sectors *N*_sec_ was computed by only counting mutant clones that were both still present at the front at the end of the simulation and that were born before *R*_f_/2. Because sectors inevitably have large areas for large s, we record the actual number of mutations *N*_mut_ in *R*_f_/2 for simulations, which was set to an average of 5 to limit interactions between clones. The scaling function saturates for ξ → 0 and scales as ξ for ξ → 0 (dashed lines are guides to the eye). The inset shows the same data on linear scale.

**Radial expansion**

To model a circular colony, one has to take into account the effect of “inflation” [5]: As the colony expands, the circumference increases in size. As a consequence, domain boundaries tend to move away from one another at a speed proportional to their current (front) distance, keeping the opening angle of the sector constant. Inflation enables mutations to fix even if they are neutral because, on long times, inflation is a stronger driving force than genetic drift. The speed 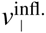 of inflation of a sector of front size *l*⊥ is such that it keeps the sector angle *l*⊥/*R* constant. Thus, we have 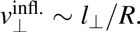 Balancing this speed of inflation with the speed 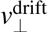 of genetic drift yields another characteristic length 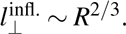 This is the establishment length for a neutral sector: If a neutral sector reaches size larger than 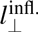, it will be protected by inflation from going extinct through genetic drift.

For the case with selection, we expect that if 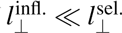, establishment will be effectively neutral as a result of the competition of drift and inflation. If on the other hand we have 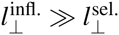 then surfing is controlled by the competition of drift and selection. This expectation can be summarized by the scaling form

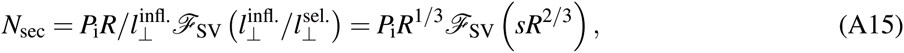
which depends on the initial radius *R* of the colony and the selective advantage *s* of the mutations. The scaling function 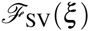 satisfies

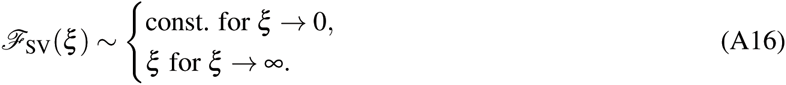

Our analysis thus predicts that when the selection coefficient is small, the number of sectors will be roughly equal to the neutral number of sectors, scaling as the third root of the initial radius. For larger selection coefficients, on the other hand, the number of sectors will scale like the radius times the selection coefficient s. This analysis is supported by simulations, see Fig. A3.

**Figure A5:**
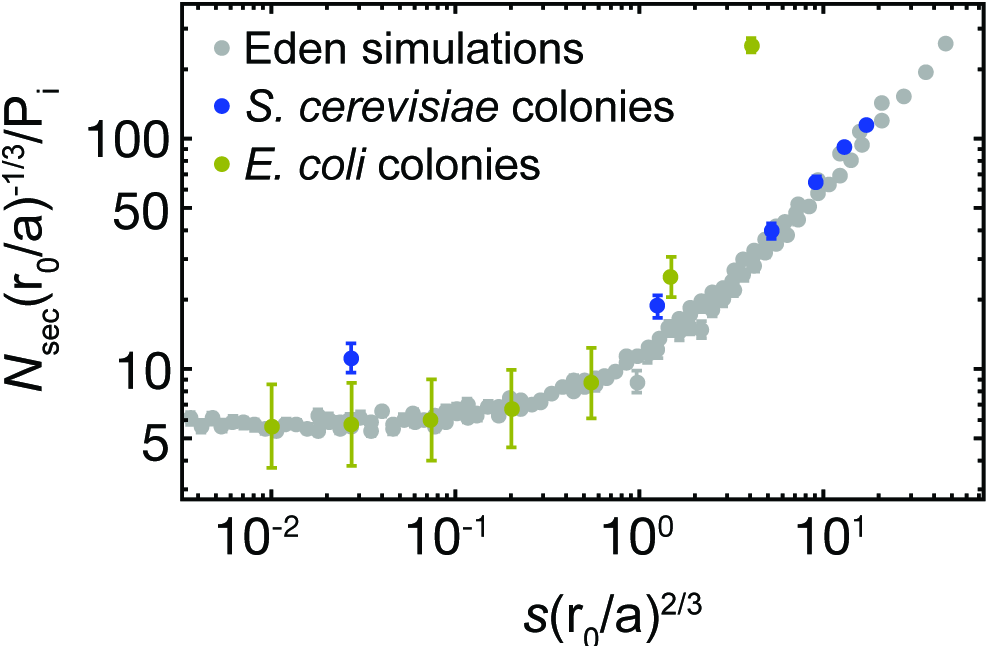
By fitting the experimental data to the scaling relation from Fig. A3, we obtain estimates for the appropriate value of *a* in our experiments. We find *a* = 0.8μm for budding yeast and *a* = 12μm for *E. coli*. Note that since the number of sectors in *E. coli* experiments depends roughly exponentially on *s*, we only fit the constant as *s* → 0. For the *E. coli* data points, we interpolated from the experimental results to obtain values for small *s*.

**3D range expansions**

The preceding discussion can be extended to the important case of three-dimensional radial range expansions, pertaining to, e.g., growing solid tumors. In 3D, a neutral surviving sector has lateral size *l*⊥ after a time of order 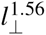 (another KPZ prediction [7]). We can estimate the surfing probability of a clone of size *l*_*c*_ by the probability *u*(*s*) ∼ (*R*/*l*_*c*_)^−2^ that a clone from a neutral mutation reaches a solid angle 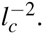 The length scale *l*_c_ again arises from the competition between drift, 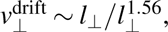 and selection, 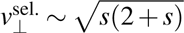 and is given by 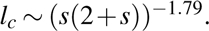 The surfing probability of a mutant with selective advantages thus scales 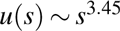 Thus, weakly beneficial mutations have a particularly small changes of surfing in three-dimensional populations.

**De novo mutations**

So far, we have focused on the number of sectors emerging from a standing variation experiment. One may alternatively consider the situation of a colony growing from a single cell. Mutations occur at a constant rate μ per lattice site. Then, we can follow very similar scaling arguments as for standing variation to arrive at the same scaling form,

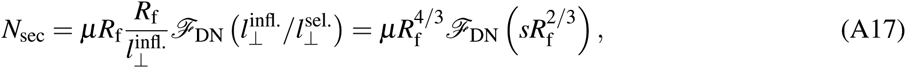
however, with a different scaling function 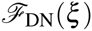 satisfying the same asymptotic limits,

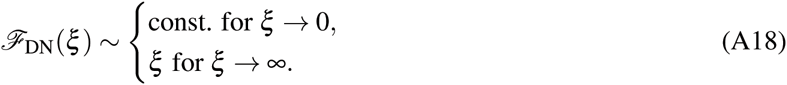

Note that the length scale R_f_ appearing in these equations defines the final radius of the colony.

**1.4 Mapping the Eden model to colonies**

The Eden model is, ultimately, a simplified lattice model that aims to capture the coarse behavior of a colony. To map Eden model predictions to an actual colony, one needs to fit the relevant phenomenological parameters.

**Figure A6:**
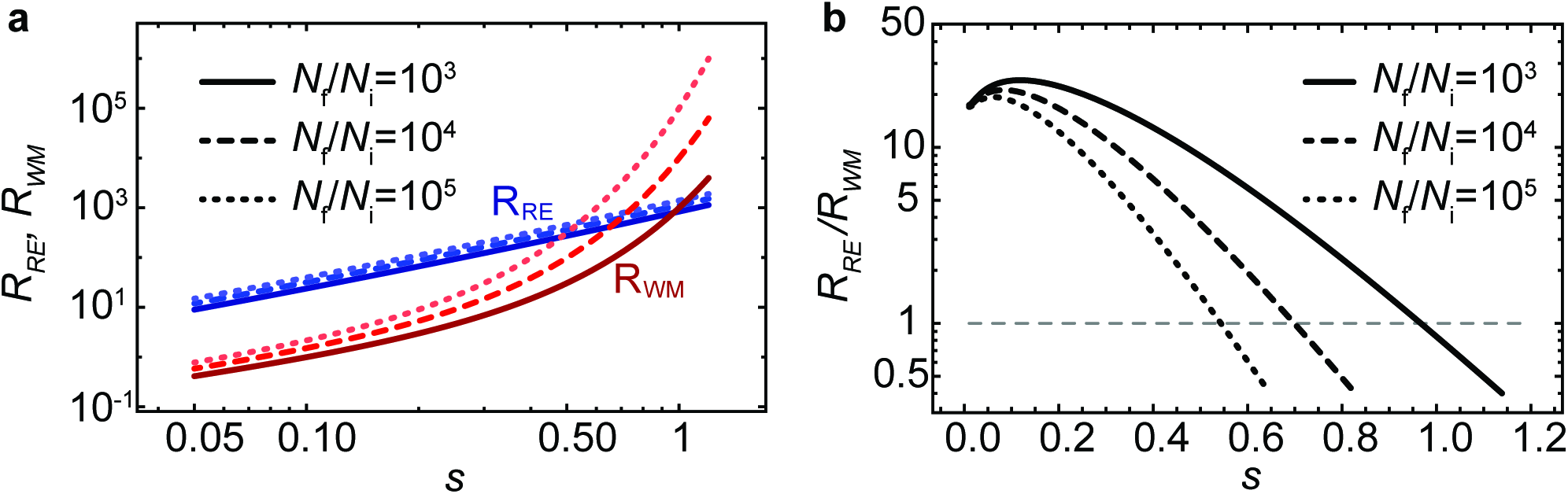
Adaptation gain in range expansions (*R*_RE_, Eq. (A12)) and uniformly grown (*R*_WM_, Eq. (A4)) populations. The number of sector was computed from Eq. (A19) with *a* = 0.8μm. A prefactor of 0.05 was introduced to match the experimentally measured sector size. We find that, for a wide range of parameters, the range expansion gives rise to a higher adaptation gain. In our experimental set-up, we expect the uniformly grown population to become more efficient than the range expansion around *s* = 0.7 (dashed line in (**b**)).

As we will see, the values of these parameters will also tell us to what extent the Eden model may be applicable.

A lattice site has a width and a length. By the rotational symmetry of a colony, we expect that we have to choose, in general different length 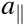 and 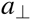 for the radial and transverse width of a lattice site, respectively. The choice of these lengths leaves selection and inflation unaffected but it influences the strength of genetic drift: Conditional on survival, a neutral sector reaches size *l*⊥, roughly, after a radial distance of order 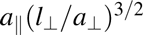, as was measured in Ref. [8]. Thus, the magnitude of the speed of growth of the width of a sector due to random genetic drift scales as 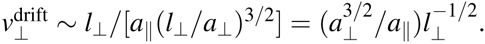 The competition between genetic drift, selection and inflation then leads to the establishment lengths 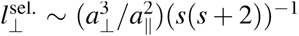 and 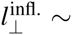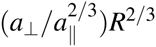, respectively. Applied to an actual colony, the Eden model prediction thus takes the form

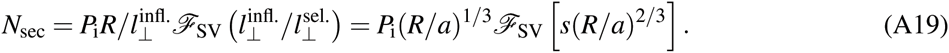

This result shows that we have effectively one parameter 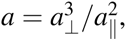, a “microscopic” length scale, to fit the predictions of the Eden model in the case of standing variation. Nevertheless, it is useful to think of this one length scale as the ratio 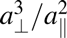 of two length scales, because there are natural candidates for the radial and transverse length scales 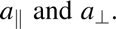 For instance, in the case of yeast, it is natural to choose the radial length to be the thickness of the growth layer and the transverse length simply as a cell diameter - there is no other transverse length scale in this problem. Then one expects 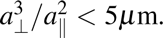 This explains then why the fitted microscopic length scale *a =* 0.8μm (see Fig. A5) is smaller than a single yeast cell diameter.

In the case of *E. coli* on the other hand, we do have another transverse length scale. Time lapse movies reveal that *E. coli* colonies buckle on length scales of order 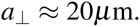 Indeed, the fitted microscopic length scale *a* = 12μm is much larger than a single *E. coli* cell.

Once the value of *a* is known, we can compare the adaptation gain in uniformly grown population and range expansions and find the parameter range for which range expansions are more efficient. Our model predicts for the case of *S. cerevisiae* (Fig. A6) and our experimental parameter range 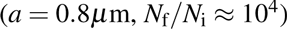 that range expansions are more efficient up to values of *s <* 0.7, although we do not expect our model to be accurate at such high values of *s*. Hence, for most experimentally accessible parameters, we expect range expansions to exhibit a higher adaptation gain than well-mixed growth.

In the case of de novo mutations, we obtain the growth layer width λ as an additional parameter, since only mutations arising at the front are able to surf. Hence,

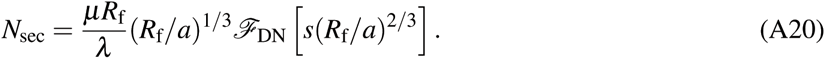

Note that, in the de novo mutation case, one has to measure both the mutation rate, growth layer width and roughness length scale to obtain predictions from the Eden model.

**Limits of the coarse-grained Eden model**

Our coarse-grained lattice model is a meta-population model, meaning that each lattice site represents a sub-population of cells. The size *N*_e_ of those subpopulations can be estimated once we have determined the linear dimensions 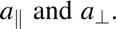 of a lattice site (see previous paragraph). Thus, we may estimate 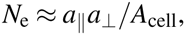 which amounts to 2.5 and 200 in the cases of budding yeast and *E. coli*, respectively.

The parameter *N*_e_ allows scrutinizing a precondition for the applicability of our coarse-grained model. If, for a given selective advantage, *N*_e_ is too large, we cannot assume that mutants will fix in a subpopulation with probability equal to their current ratio. This is assumed when we set the mutation rate in the Eden model equal to the mutation rate of single cells. The same assumption is made in the case of standing variation, when we assume that the initial fraction of mutant lattice sites is equal to the initial frequency of mutant cells. If subpopulations behaved like well-mixed sub-populations, for instance, we would have to require 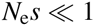. Note that this condition is strongly violated in our *E. coli* experiments. For 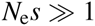, the effective mutation rates as well as the initial frequencies would have to be multiplied by *N*_e_s. Since, however, our populations are manifestly spatial, it is not clear how a more microscopic model would behave. Therefore, we also implemented more explicit simulations that take into account the shape and steric interaction between cells (described in detail below).

**1.5 Analysis of long-range jumps**

We extend our meta-population by allowing for long-range jumps in each step. The rationale behind introducing long-range dispersal is that well-mixed growth and colonies are natural opposites in that they feature no and strong spatial correlations and mixing. Long-range jumps allow for a breaking of spatial correlations and thus lie in between these two cases. We then also expect the adaptation efficacy to interpolate between the colony and the well-mixed case as the likelihood of long-range jumps increases.

As described in the Methods section above, we can vary the likelihood of large jumps by tuning the parameter *JJL*. Hallatschek and Fisher [1] showed that the expansion speed of a growing population, in terms of its range, depends on the parameter *δ* = *μ* — *d*, where *d* is the spatial dimension. If *δ* > 0, the range of the population grows as a power-law, while for *δ* < 0, long-range jumps are frequent and the range of the population grows as a stretched exponential.

Introducing mutants with selective advantage *s* at initial frequency *P*_i_ = 0.02, we studied the influence of dispersion range on the efficiency of adaptation. The naive expectation would be that in the limit of short-ranged jumps, adaptation should be as efficient as in the classical Eden model, whereas long-range jumps increase the mixing of the population such that adaptation becomes less efficient, asymptotically becoming well-mixed.

Fig. A7 shows examples of populations grown from *N*_*i*_ = 10^3^ to *N*_f_ = 10^6^ for different values of *μ* and *s*. We observe that the final frequency of the mutants increases with *s*, as expected, but does so much more strongly when *μ* is large. As *μ* increases, the populations become increasingly patchy, with mutants primarily residing in confined spatial regions. For *μ* = 5, we even observe sectors very much like in Eden simulations. Fig. A8 shows the results of 750 simulations for each set of parameters (*μ*, *s*). We see indeed that the average final frequency of mutants increases as long-range jumps become increasingly rare. Thus, long-range jumps can hinder adaptation from standing variation even in spatially structured growing population because they effectively induce mixing which allows previously trapped clones to continue to grow, allowing for fewer generation to happen at the very front (for fixed final population size).

**Figure A7:**
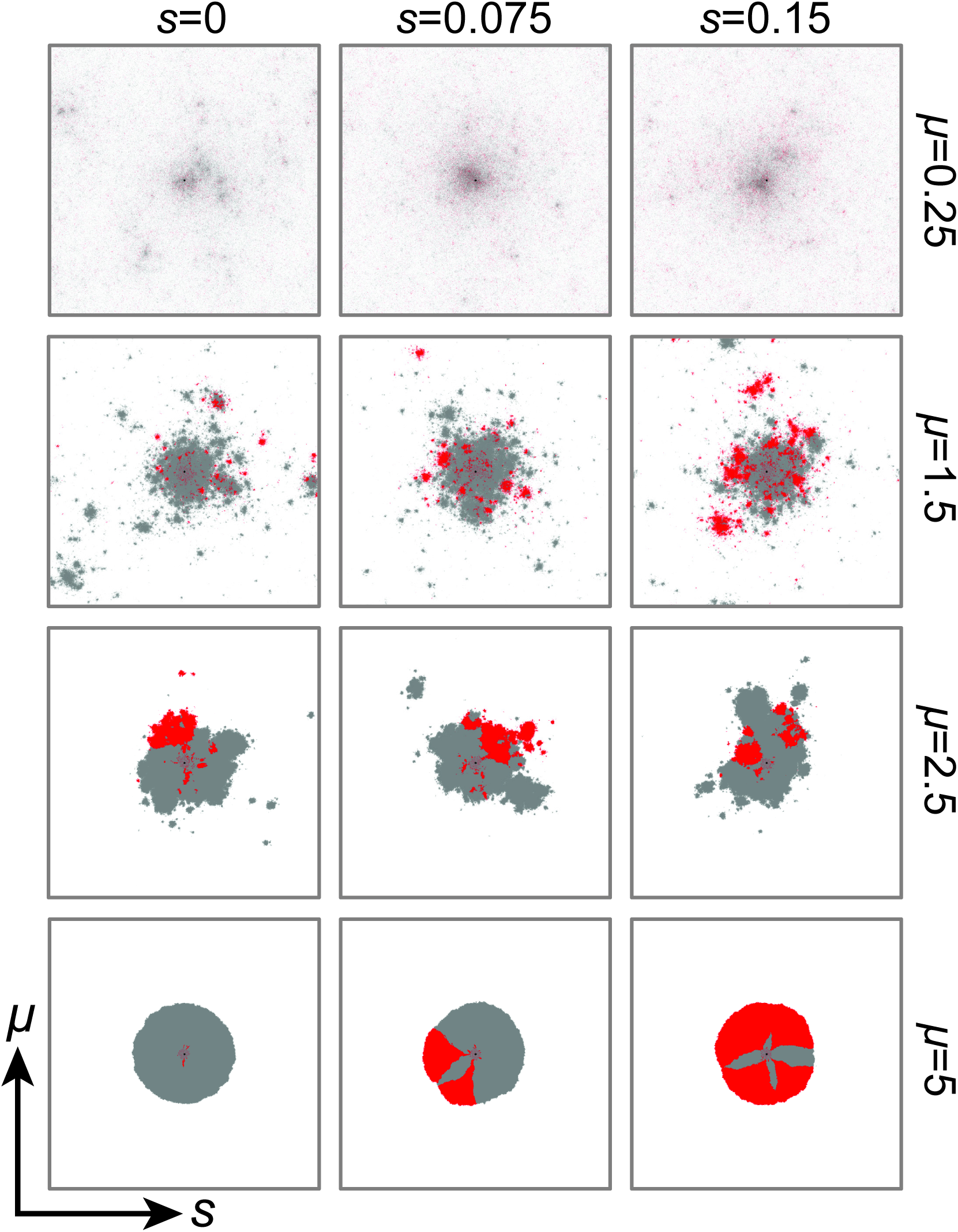
Examples of populations undergoing range expansions with long-range dispersal. Different selective advantages s of the mutants (shown in red) over the wild type (gray) are shown as the columns. Varying the “spread” coefficient μ (rows), we obtain almost well-mixed populations for small μ (when large jumps are common), while sectors emerge for very large μ (when practically no large jumps occur). For intermediate μ, mutants accumulate in patches that resemble sectors more and more as μ increases. The larger μ, the higher the mean final mutant frequency, as shown in Fig. A8.

**Fig. A8.**
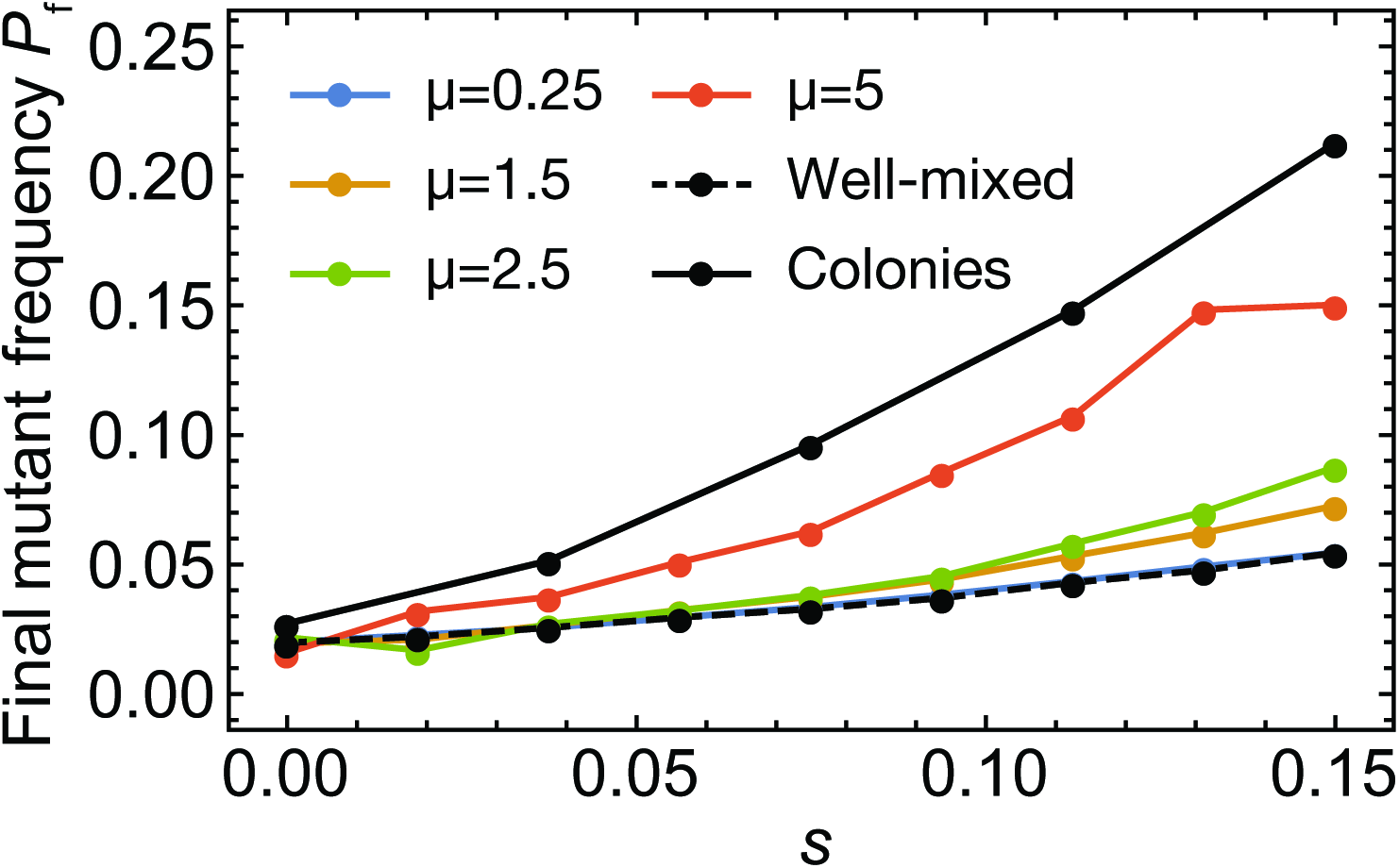
Final mutant frequency in populations grown with varying degrees of dispersal. All populations grown from *N*_*i*_ = 10^3^ to *N*_f_ = 10^6^ with a starting fraction of *P*_i_ = 0.02. Populations with long-range jumps show adaptation efficiency intermediate between well-mixed and strictly short-ranged range expansions (Eden model). For μ ≪ *δ*, the final mutant frequency becomes indistinguishable from the well-mixed case. Results obtained by averaging over 750 simulations.

**Table A1:** Parameters of the off-lattice model.

| Parameter | *E. coli*-like (long) cells | Yeast-like(short) cells |
| --- | --- | --- |
| Diffusion constant $D$ [$\mu\text{m}^2/\text{h}$] | 500 | 500 |
| Initial nutrient concentration [a.u.] | 1 | 1 |
| Nutrient uptake rate $\kappa$ [a.u./h] | 1.5 or 1.8 | 4 |
| Young modulus $E$ [kPa] | 100 | 500 |
| Minimal division time $T$ [min] | 40 | 120 |
| Cell diameter [$\mu\text{m}$] | 1 | 4 |
| Minimal length [$\mu\text{m}$] | 2.5 | 0 |
| Damping coefficient $\zeta$ [$10^{16} \text{ h}^{-1}$] | 65 | 65 |
| Friction coefficients | $k_{\perp} = 4, k_{\parallel} = 0.25$ | $k_{\perp} = k_{\parallel} = 1$ |
| Width of simulation box $L$ [$\mu\text{m}$] | 340, 640 | 320, 640, 1280 |

**2 Individual-based simulations: method and results**

**2.1 Model description**

In order to develop a microscopic understanding of the surfing process, we used a model based on that used in Ref. [2], with a few modifications. Our model strikes a balance between computational cost, limited knowledge of the nature of mechanical interactions between cells in microbial colonies, and the reproduction of experimental observations made in this work.

All cells are modeled as sphero-cylinders of variable length *l* and identical radius *r*_0_. Cells interact mechanically through Hertzian repulsion: 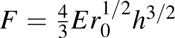 where *F* is the repulsive force, *E* is the effective Young modulus of the cell, r_0_ is the radius and *h* is the overlap between interacting cells. The dynamics of the cells is described by the overdamped Newton equations of motion:

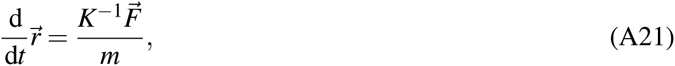

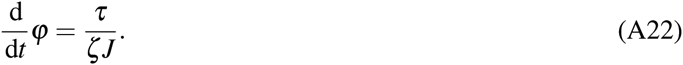

Here, 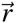 is the position of the cell’s center of mass, *φ* is the angle the cell with the x-axis, 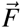 is the total force, *τ* is the total torque acting on the cell, *m* is the mass, *J* is the momentum of inertia of the sphero-cylinder, and *ζ*is the damping (friction) coefficient. The matrix *K*

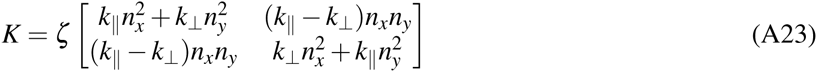
takes into account the possible anisotropy of friction between the cell and the surface: *k*_⊥_ is the damping coefficient in the direction perpendicular to cell’s major axis, 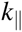 is the damping coefficient in the parallel direction, and 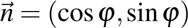. For isotropic friction, 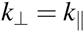, and the matrix *K* reduces to the identity matrix times the friction coefficient. To model cells which prefer to roll rather than to slide we set 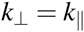, whereas for cells that prefer to slide along the major axis it holds that 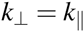. In particular, for “yeast-like” cells we assume isotropic friction, whereas for *”E. coli-like”* cells we set 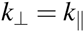. This replicates experimentally observed long “chains” of aligned cells and high surface roughness of *E. coli* colonies.

**Table A2:** Steady-state properties of the growing layer.

| Quantity | *E. coli*-like (long) cells |  | Yeast-like(short) cells |
| --- | --- | --- | --- |
| | $\kappa = 1.5$ | $\kappa = 1.8$ | $\kappa = 4$ |
| Thickness of the growing layer $\lambda$ [$\mu\text{m}$] | 31 | 25 | 95 |
| Roughness of the growing layer $\sigma$ [$\mu\text{m}$] | 12 | 15 | 3.6 |
| Speed of the growing layer [$\mu\text{m}/\text{h}$] | 26 | 23 | 31 |
| Average area $\langle A \rangle$ per cell [$\mu\text{m}^2$] | 1.7 | 2.0 | 15.6 |
| Average linear size $a$ of cell [$\mu\text{m}$] | 1.3 | 1.4 | 4.5 |
| Thickness of the growing layer $\lambda/a$ [cells] | 24 | 18 | 24 |

Cells consume nutrients diffusing in the 2D substrate beneath the colony of cells. The concentration 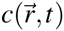 of the nutrient evolves in time as

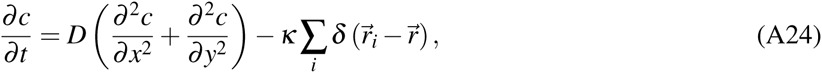
where *D* is the diffusion constant, icis the nutrient uptake rate and 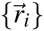 are the positions of the cells. We assume that cells elongate at a constant rate as long as the local nutrient concentration is larger than 2% of the initial concentration, and divide when they double in length. The length of individual cells thus increases linearly in time in our model. Although this is not true for real microorganisms [3, 4], deviations from linear growth are not important for the population level we are concerned with.

We model faster-growing mutants by increasing both the elongation rate and the nutrient uptake rate by 1 + *s*, where *s* is the selective advantage of the mutant over the wild type.

To reduce computation time we simulate only a narrow strip of width *L* at the front of the colony, with periodic boundary conditions in the direction perpendicular to the direction of growth, and fix cells which lag behind the growth layer.

All parameters are listed in Table A1. The assumed values have been chosen to make simulations computationally feasible while at the same time to approximately reproduce experimental observables: the average cell size, the velocity of the moving front, and the thickness of the growth layer. For example, the trade-off between speed and realism required the diffusion constant to take an unrealistically small value.

**2.2 Characterization of the properties of simulated colonies**

We define the growth layer as the layer at the colony front in which cells were replicating. We calculated the thickness *λ* of the growth layer as the average of the shortest distances between cells at the very front of the growth layer (first line of cells) and the last layer of cells towards the bulk still exhibiting growth.

The roughness *σ* was defined as the square root of the mean square deviation of the front height *y*(*x*), where *y*(*x*) corresponds to the envelope of the front, with resolution 1*μ*m. The speed of the front was obtained by fitting a straight line to the average position of the front *ӯ*(*t*).

Fig. A9 shows that the thickness and the roughness of the growing layer stabilize after some time. The steady-state values are given in Table A2. The table also shows the average cell size determined as the area of the growing layer divided by the number of cells. This is the actual size taken by the average cell; mechanical compression due to growth causes this area to be slightly lower than the average area of an isolated spherocylinder as determined by the parameters from Table A1.

**Figure A9:**
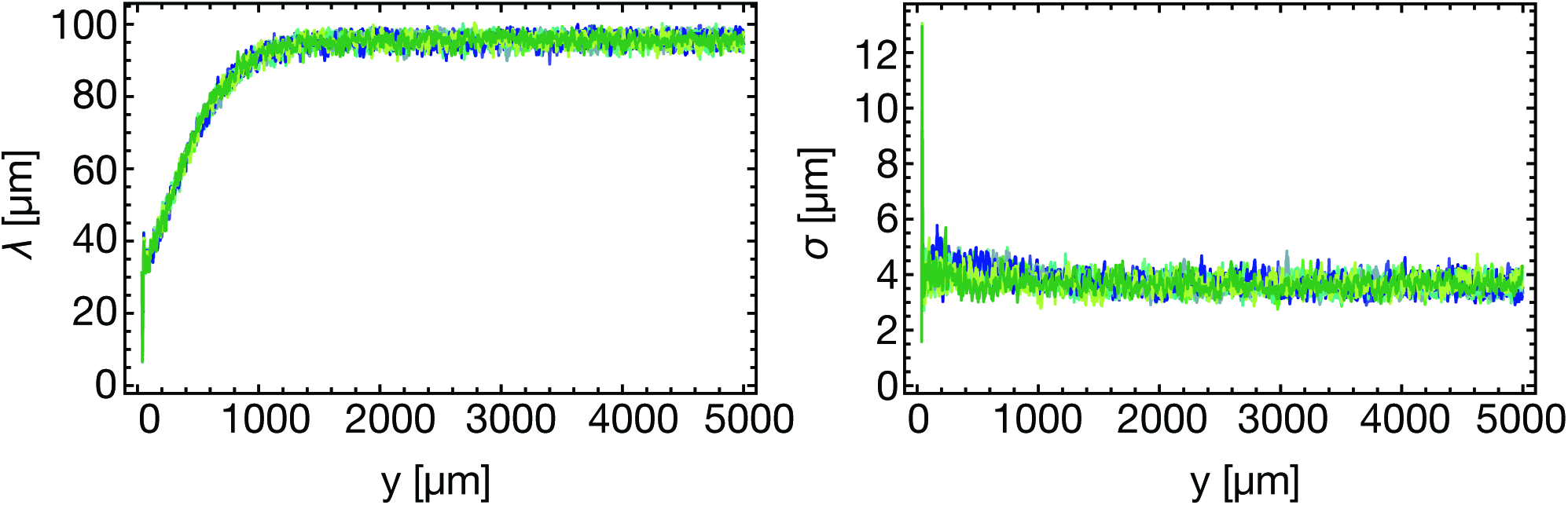
Run-in period of off-lattice simulations. Thickness *λ* and roughness *σ* of the growing layer for “yeast-like” cells, for 10 simulation runs (different colors).

**Figure A10:**
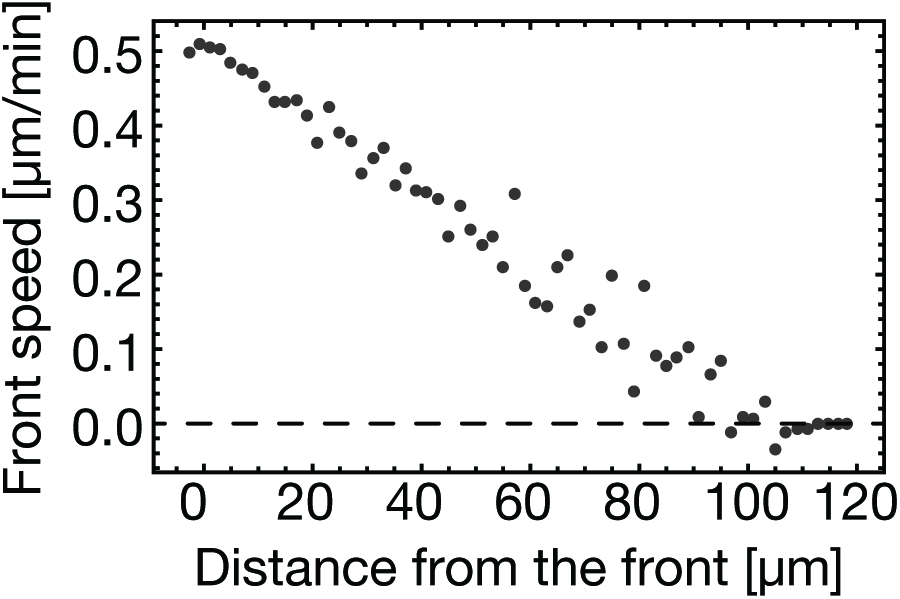
Average speed of cells at a given distance from the colony front for simulated “yeast-like” cells.

We also computed the average linear cell size *a* as the square root of the average area, *a* = 〈A〉^1/2^. This enabled us to express the thickness of the growing layer in cell lengths as *λ/a*. We adjusted the parameters of the model for “yeast-like” and *”E. coli-like”* cells such that *λ/a* was approximately the same for both types of cells.

The speed of the cells in the growing layer is a linear function of the distance from the front (Fig. A10). This replicates well the experimentally observed behavior (Fig. SIE 8). We note that in our experiments cessation of growth in the center of the colony and the emergence of the growing layer may be due to the accumulation of waste rather than nutrient exhaustion. However, as demonstrated in Ref. [2], the behavior of the model is similar regardless of whether growth is limited by nutrients or waste products, and that in both cases growth becomes confined to a thin layer after an initial period of exponential growth, in agreement with what is observed experimentally.

**Figure A11:**
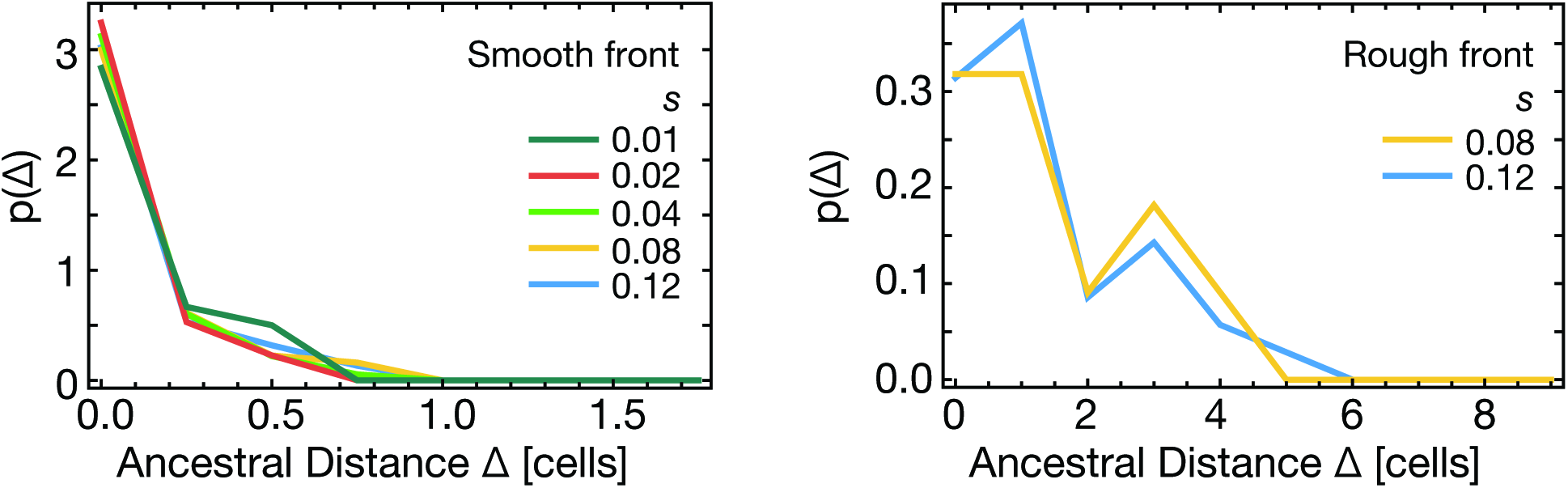
The probability density that a lineage originated at distance Δ from the front (cell lengths) given that it fixed in the growing layer, for different selective advantages (s = 1%, 2%, 4%, 8% and 12%). Left: “yeast-like” cells. The probability is concentrated at the very first line of cells, almost independently of selective advantage. Right: “E. coli-like” cells. The distribution is slightly broader but still concentrated around Δ = 0.

**2.3 Surfing probability at the front and distribution of ancestor location**

To determine the surfing probability *P*_surf_ of mutants with different selective advantages we first ran simulations in which mutant cells were randomly inserted into a steady-state growing layer. We ran between 1000 and 10000 simulations and calculated *P*_surf_ as the proportion of runs in which the mutant fixed in the growing layer. We also determined *P*_surf_ for mutants appearing at different distances from the front.

Our results show that *P*_surf_ is very small even for quite large selective advantage s = 0.12: *P*_surf_ = 0.004 for “E. coli-like” cells and *P*_surf_ = 0.015 for “yeasts-like” cells for parameters as in Table A1. Fig. A11 shows that the surfing probability quickly decreases with the distance Δ from the front of the first (founder) mutant cell.

We then ran simulations with mutants inserted only in the first line of cells. Fig. A12 shows that “yeast-like” cells have a much larger *P*_surf_ than “E. coli”-like cells. Since the two cases differ in the roughness of the growing layer (c.f. Table A2), we hypothesized that roughness is the main factor affecting the amount of genetic drift. To test this, we simulated “E. coli”-like populations with reduced roughness – this was achieved by decreasing the nutrient uptake rate (Table A2). We indeed observed an increase in *P*_surf_, in accordance with our hypothesis.

**3 Supplementary discussion: Dynamics behind the front**

In our experiments, change in local allele frequencies occurs only directly at the front, and our analysis above reflects this fact. While true for non-motile microbes, our arguments arguably extend to other cases where spatial arrangements are mostly conserved, e.g., biofilms and to some extent, solid tumors. However, our results are valid more generally, independently of whether sectoring is neutral or beneficial, as long as the front advances faster than the blurring of the sectors occurs. However, if there is mixing behind the front, any spatial inhomogeneity in local allele frequencies will eventually be blurred out.

**Blurring of neutral sectors**

If individuals can move randomly behind the front, existing sector boundaries will undergo diffusion and thus have a characteristic width *w* scaling as 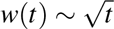. The front, however, advances at constant speed and hence the front position *r*(*t*) ∼ *t*. Hence, on long time-scales, the advancement of the front is much faster than the blurring of sectors, and sector boundaries will remain sharp near the advancing front.

**Figure A12:**
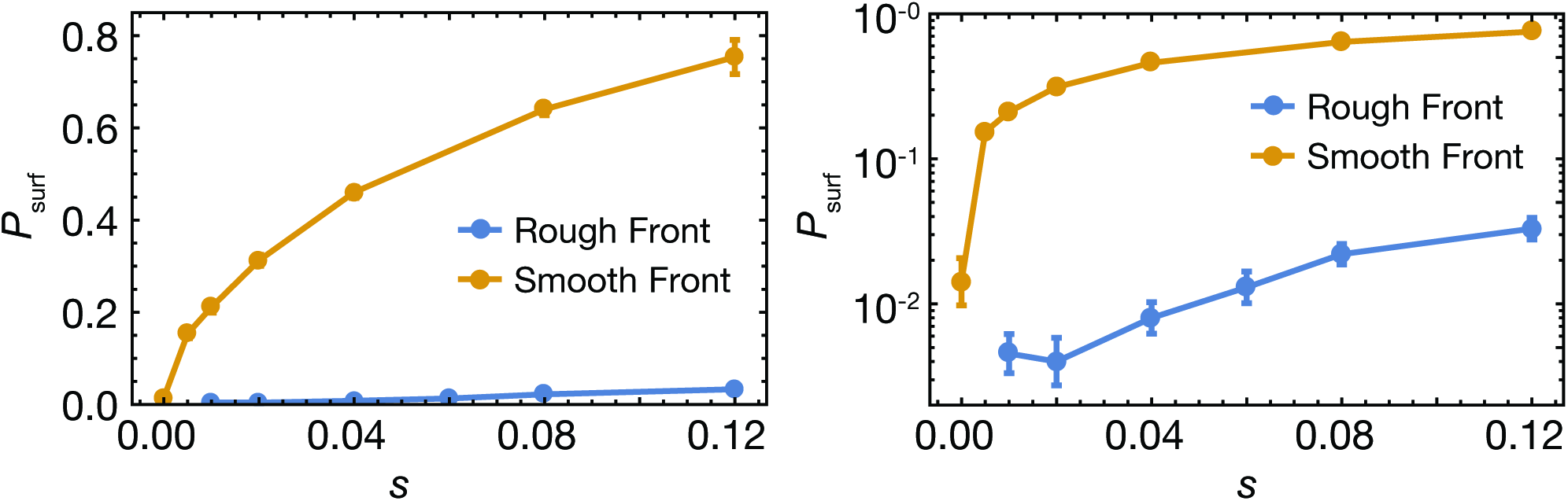
Left: Surfing probability *P*_surf_ versus selective advantage s for “yeast-like” cells (“smooth front”, orange) and “E. coli”-like cells (“rough front”, blue. Here, *κ* = 1.8). Smooth fronts deviate from a line only by about 1 cell diameter, while rough fronts exhibit a roughness of about 10 cell diameters. Parameters and measured characteristics of the populations are given in Tables A1 and A2. Mutants were introduced only into the first layer of cells. The surfing probability *P*_surf_ decreases with increasing roughness, but increases with selective advantage. Right: the same plot with a logarithmic scale.

**Beneficial sectors behind the front**

After the front has passed, beneficial sector will slowly blur due to diffusion, but may also widen or shrink as the beneficial mutants compete with the wild type in the bulk. Even if the mutants exhibit a growth rate advantage at the front, there is not a priori reason to assume that the same is true in the bulk, where other characteristics than maximum growth rate may be more important. For example, a more efficient use of nutrients in poor growth environments (like the bulk of a colony) may prove to be more advantageous than the ability to outgrow one’s competitors when nutrients are abundant.

## Appendix B: Additional experimental results

**Figure B1:**
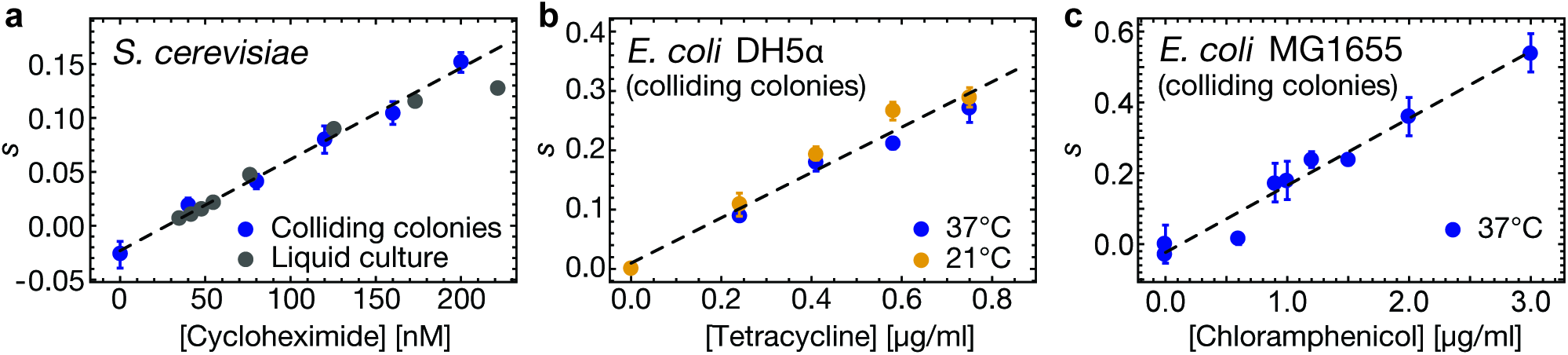
Selective advantages *s* between resistant and sensitive strains as function of drug concentration for *S. cerevisiae* and *E. coli* for different assays and conditions. (a) Budding yeast strains with W303 background (yMM9 and yJHK111) used in Fig. 1. Best linear fit is shown and used throughout the paper. Liquid culture fitness measurements (3 replicates from the same initial culture per data point, gray dots) over 60 generations agree with the colliding colony result (blue dots) for a range of cycloheximide concentrations. (b) *E. coli* DH5α competition (strains eOH2 and eOH3) on plates with different concentrations of tetracycline at 37°C (blue data points) and 21°C (yellow data points) using the colliding colony assay. Temperature had no significant impact on the relative fitness of the two strains. Best fit is shown through combined data and used throughout the paper. (c) *E. coli* strain MG1655 competed against strain SJ102 at different concentrations of chloramphenicol, measured using the colliding colony assay, with linear best fit. All error bars are standard error of the mean (about 20 replicates per data condition, except for well-mixed competition).

**Figure B2:**
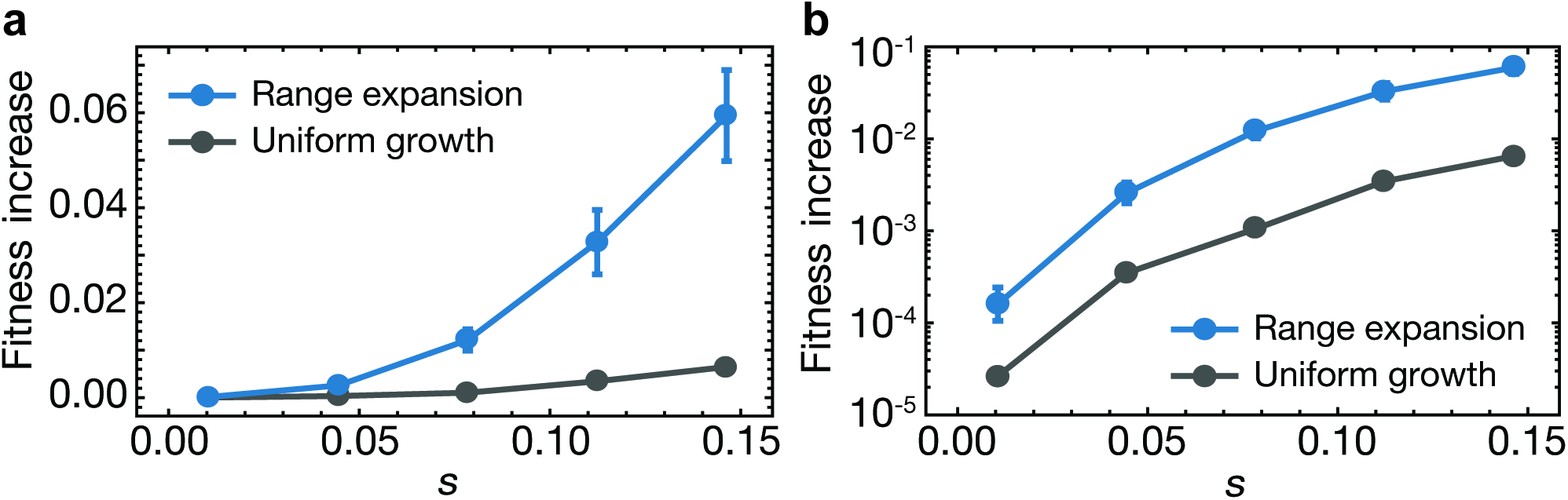
Increase in mean fitness, defined as 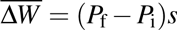, computed from the final mutant frequency *P*_*ƒ*_ measured in Fig. 1h, with linear (a) and logarithmic axis (b). Range expansions leads to a higher increase in mean fitness than uniform growth to the same final population size.

**Figure B3:**
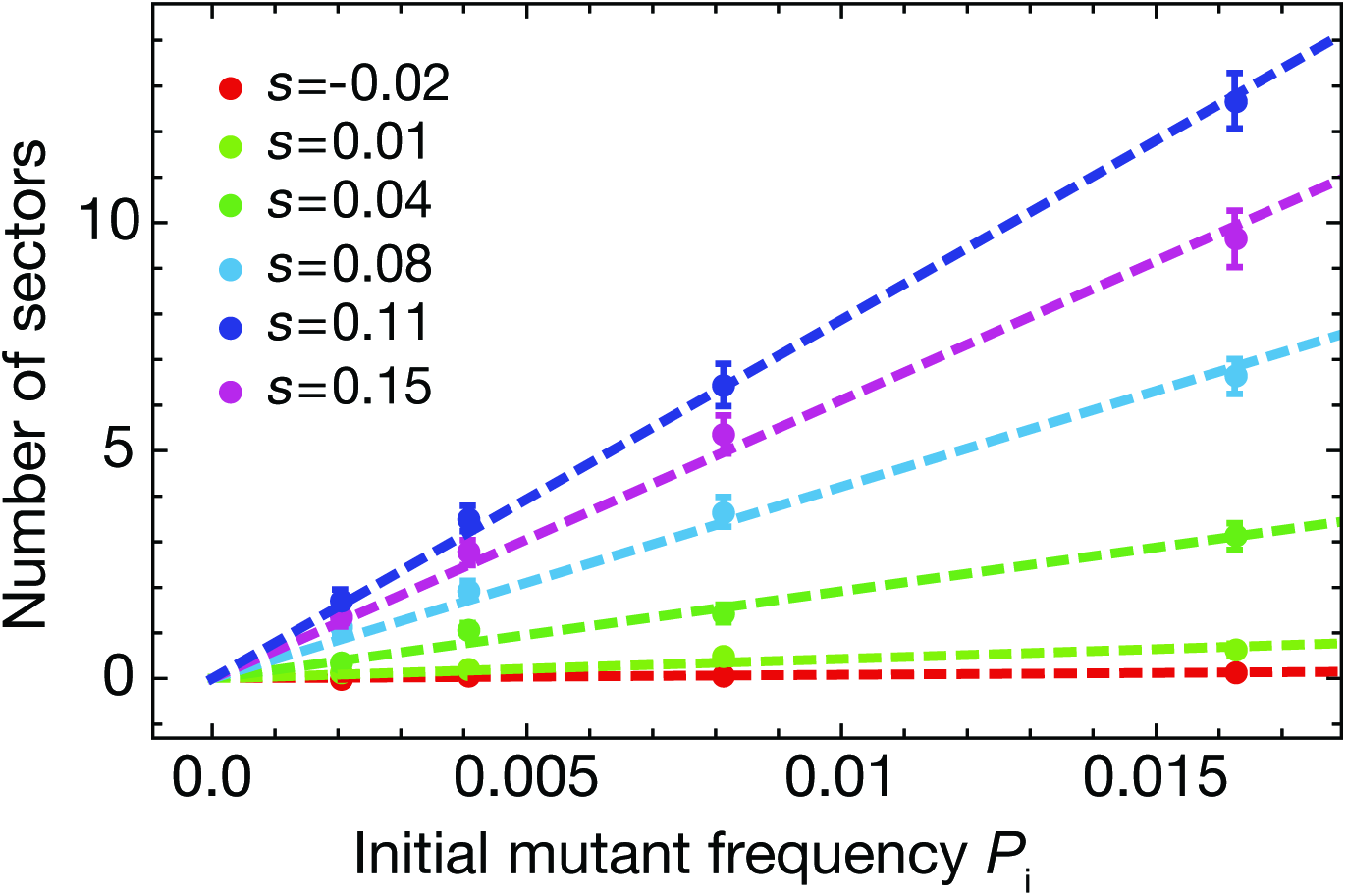
Number of sectors *N*_sec_ counted in yeast colonies at different initial mutant frequencies *P*_i_ and selective advantages *s*. The proportionality of sector number and initial mutant frequency implies that sectors arise independently for small enough fractions. Points are averages from 30 colonies per condition, error bars correspond to the standard error of the mean.

**Figure B4:**
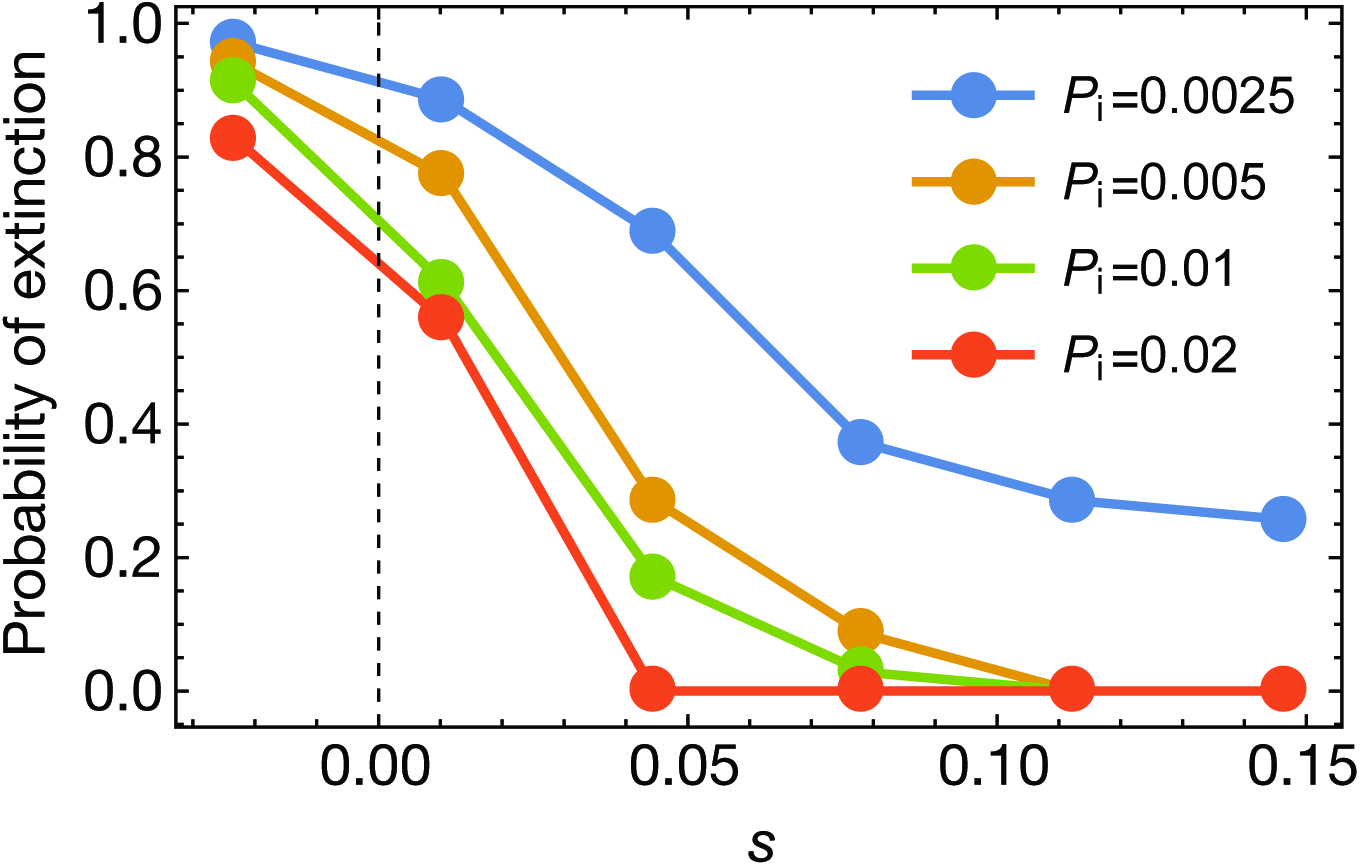
Probability of extinction, defined as the probability of having zero sectors at the front, in yeast colonies for a variety of initial mutant frequencies *P*_i_ and selective advantages *s* (35 colonies from same initial culture per data point).

**Figure B5:**
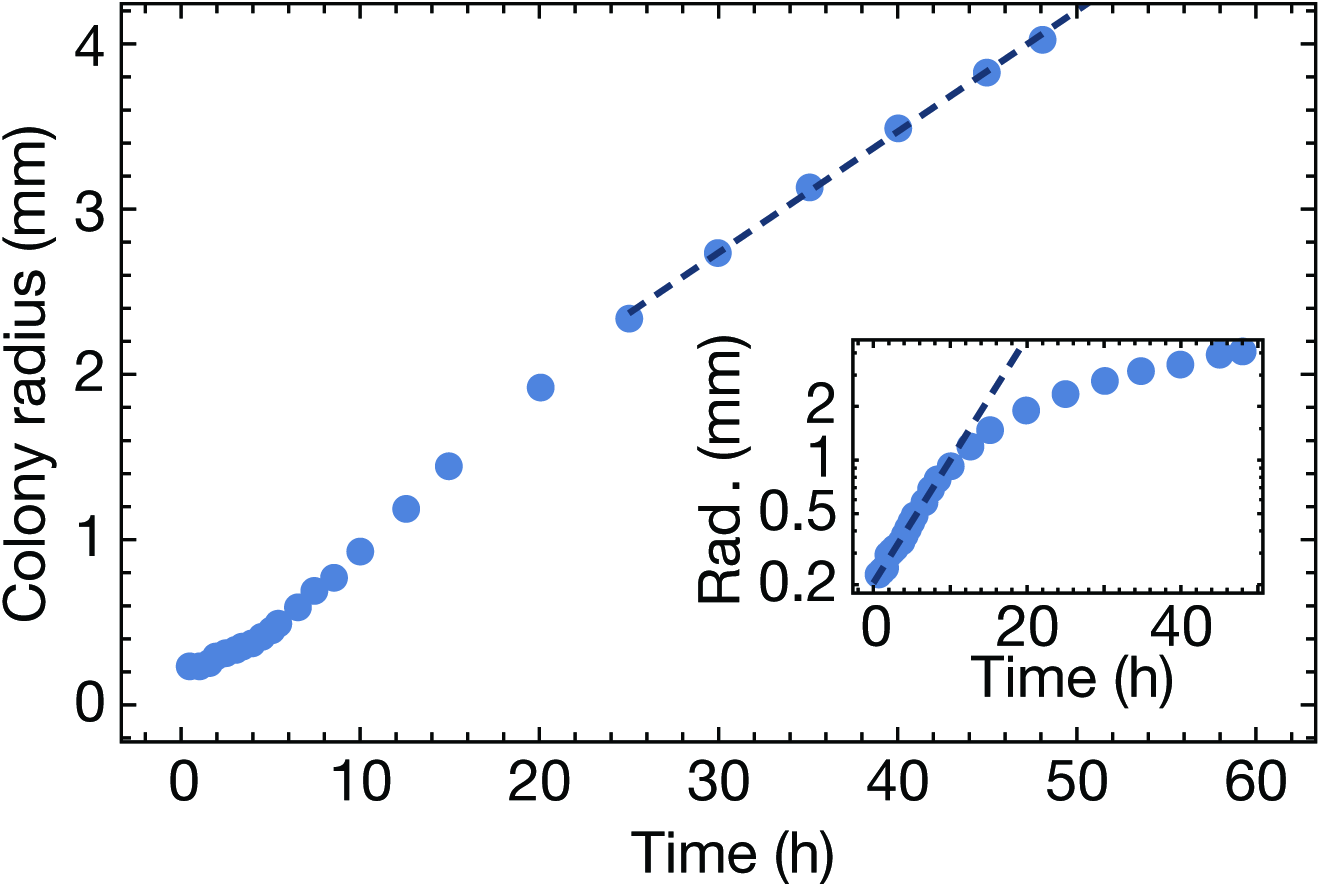
Two regimes of yeast colony radius growth. Colony radii extracted from a time lapse movie of a growing yeast colony. Single yeast cells were inoculated onto an agar plate and grown for about 12 hours. Once microcolonies were detectable, the agar plate was transferred to a stage-top incubator and the colony was imaged in bright-field and fluorescence light every 30 minutes. Initially, the colony radius growth exponentially, indicating that the radius of the colony is not yet larger than the eventual growth layer thickness of the colony. For late times, the colony radius grows linearly in time, which can be interpreted as a growth layer of constant final thickness.

**Figure B6:**
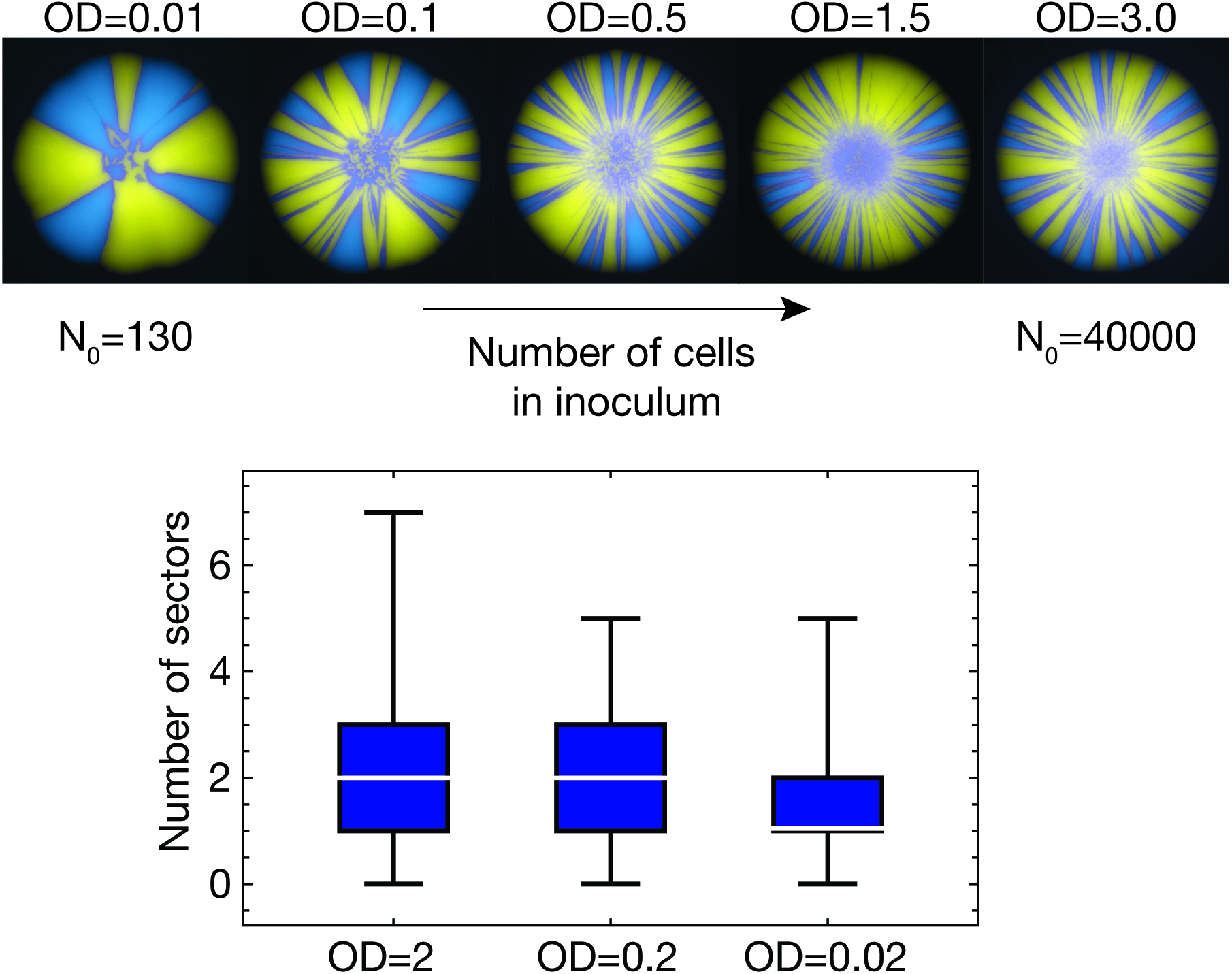
Number of sectors for different number of cells in inoculum. Top: Example images show the change of population patterning with increasing cell number in the inoculum. Here, we mixed two selectively neutral S. cerevisiae strains (yJHK111 and yJHK112) at an initial ratio of *P*_i_ = 0.5. For large cell number *N*_0_ the population pattern does not change when increasing N_0_ further. (Bottom) Number of sectors measured in standing variation assays for different inocula, for the same strains. We assayed 50 neutral colonies with an initial ratio of *P*_i_ = 0.01 per dilution, allowing us to count individual sectors. We found no significant difference in sector numbers when diluting a typical culture (OD=2 in the figure, corresponding to about 30000 cells in a 2μl droplet) by a factor of 10 (*p* > 0.05, Mann-Whitney U-test). Dilution by another factor of 10 showed again no significant difference to the intermediate case (first 10-fold dilution), but did show a significant difference (*p* < 0.05) to the original case. Taken together we conclude that the number of sectors is not sensitive to density of the initial culture, given that the inoculum contains at least about 1000 cells in a 2μl droplet. This means that typical pipetting errors or a small change in cell densities of the culture mixtures, which we estimate to be in the range of at most 10 per cent, should have no impact on the number of sectors. The observations can be understood from Fig. 3: Only cells at the front have a chance to surf, and in our experiments, the front is imposed as an initial condition by the drying inoculum. Hence, as long as the cells in the droplet form a continuous ring of cells (such that at every point of the ring there is a defined front), the number of cells in the bulk of the inoculum plays no role in the future fate of the colony.

**Figure B7:**
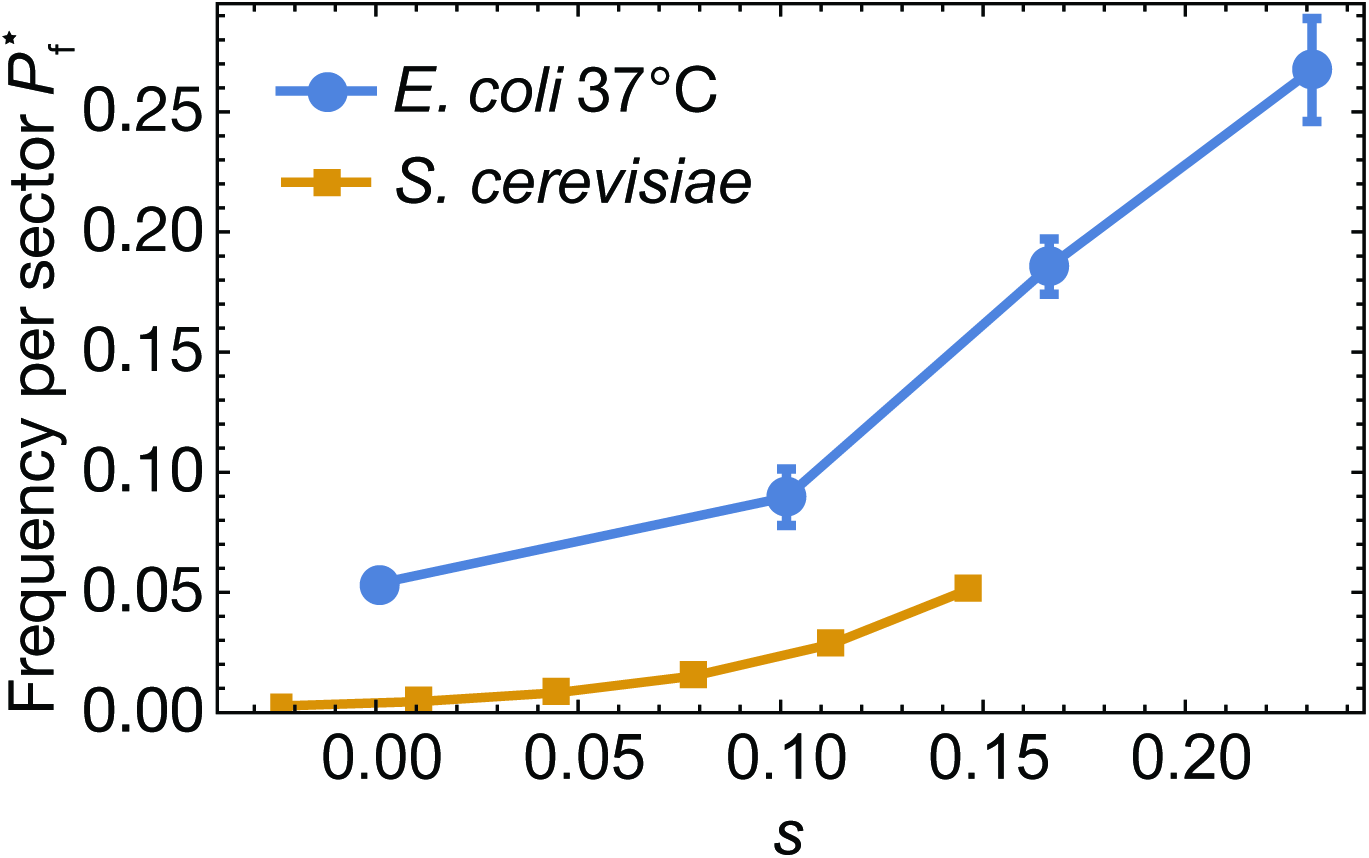
Frequency per sector for *E. coli* DH5α grown at 37^°^C (blue dots, see also Fig. 4f) together with corresponding data for *S. cerevisiae* (data as in Fig. 1j). Only individual, non-interacting sectors were selected for analysis. Each individual sector is much larger in (relative) size in *E. coli* than in *S. cerevisiae* colonies. In fact, a yeast sector at the highest assayed selective advantage has a relative size comparable to a neutral *E. coli* sector. Error bars are standard errors of the mean, from about 20 sectors per selective advantage *s*.

**Figure B8:**
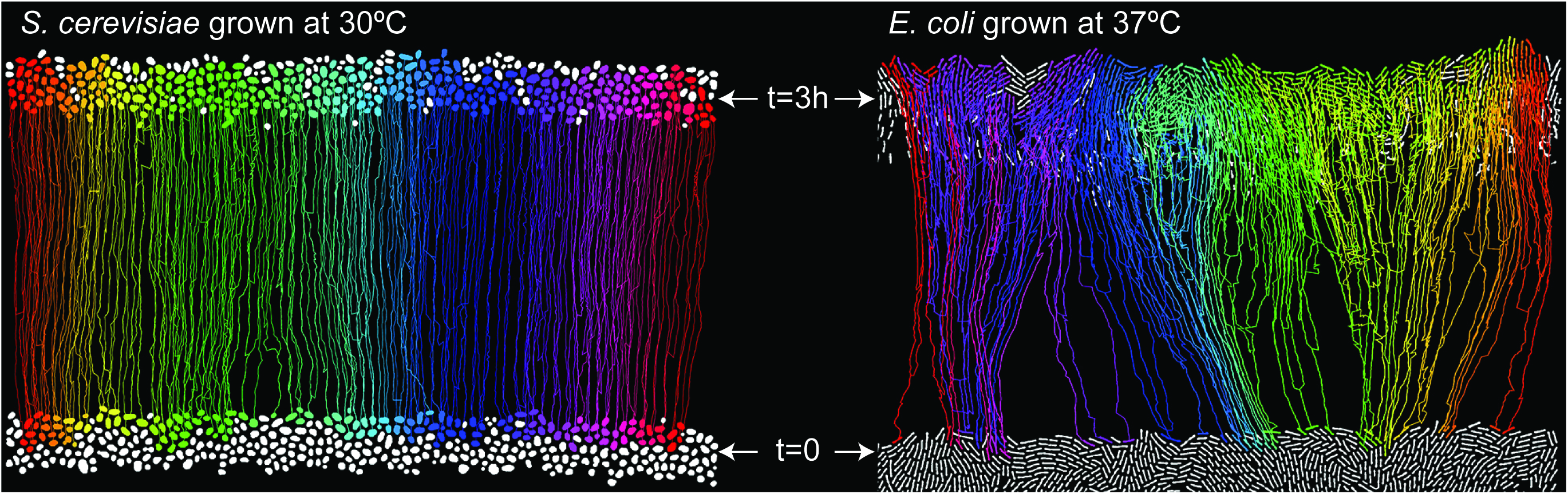
Tracking of individual cell dynamics in *S. cerevisiae* and *E. coli* front reveal microscopic motion patterns. *S. cerevisiae* and *E. coli* fronts tracked for 3 hours (full images from Fig. 3). Shown are the front of each colony at the start of the experiment (*t*=0) and after 3 hours. All tracked lineages and cells are colored. White cells at the top could not be tracked, while white cells at the bottom have no tracked descendants at the front after 3 hours.

**Figure B9:**
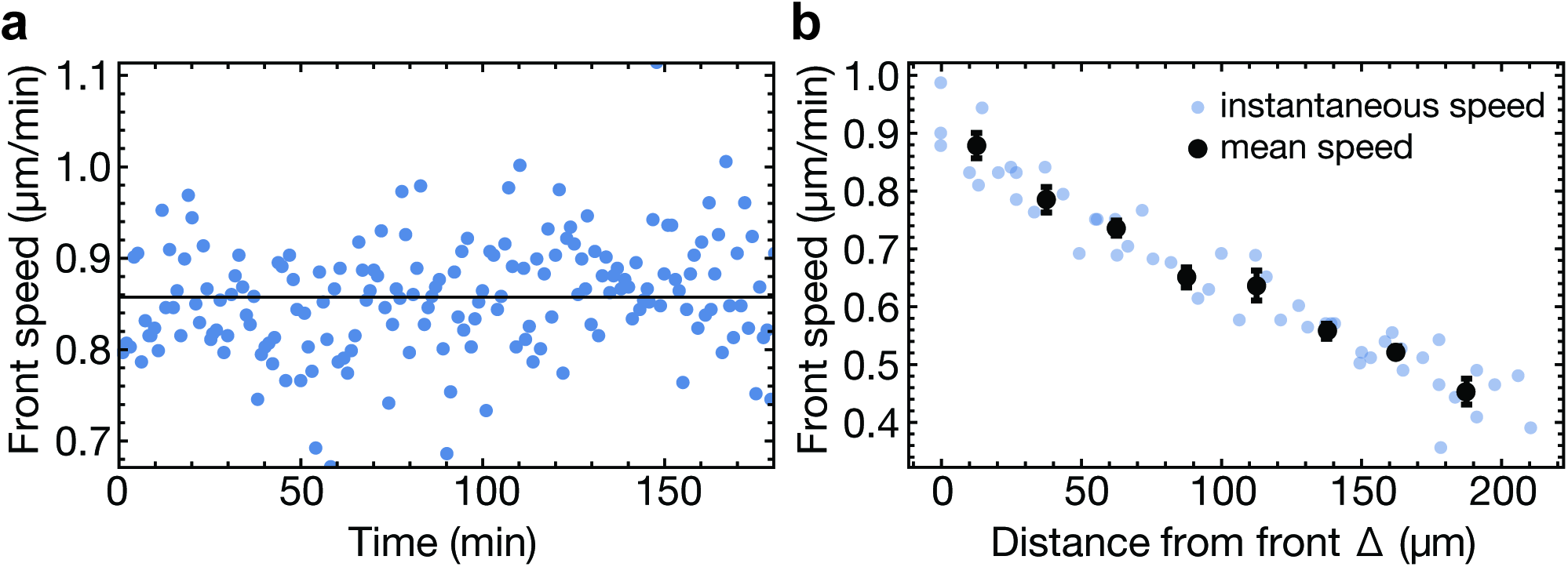
Overall front speed and speed of individual cells at and behind the front for *S. cerevisiae*. A. Instantaneous front speed, defined as the difference in mean front position in each frame, and time average (solid line). The front speed in a *S. cerevisiae* colony is constant over at least three hours (from SI movie 1, see also Fig. B5). B. Speed of individual cells as a function of distance from the front. Speed was measured by visually following 16 individual cells, initially a distance Δ behind the front, and recording their position every 30 minutes for 90 minutes. Instantaneous speed was computed by dividing the relative change in position by 30 minutes. Here, Δ is the distance from the front at the beginning of each 30 minute interval. For the mean speed, data were binned in 25μm intervals. An approximately linear decrease in speed implies near-constant growth rate at least 200μm into the bulk of the colony.

**Figure B10:**
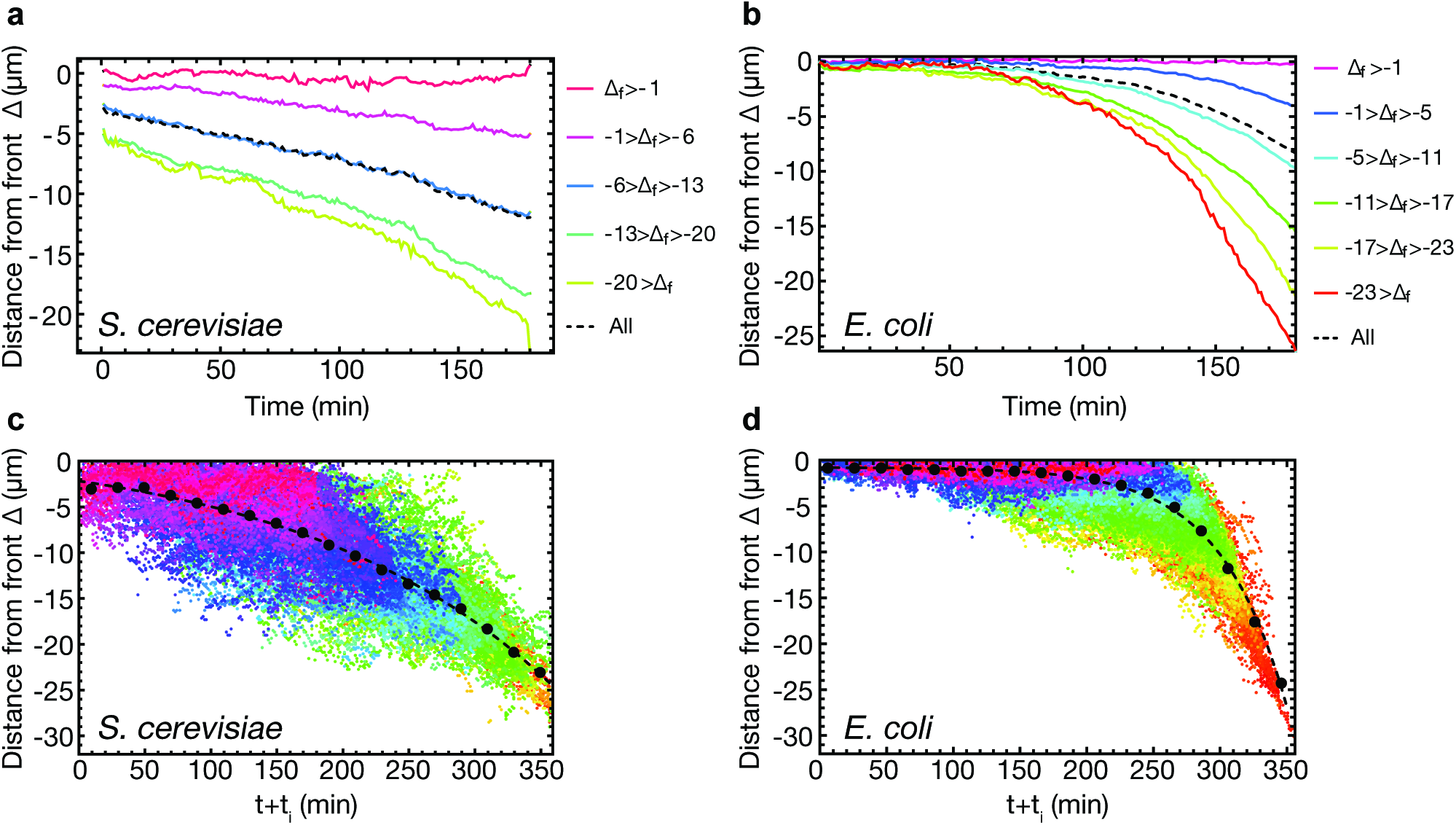
Dynamics of lineages at and behind the front, extracted from SI movies 1 and 2, for *S*. cerevisiae and *E. coli*. Front position was recorded at every time point, and the distance to the front computed for all cells. For (a) & (b), cell trajectories were pooled together depending on their final distance Δ_f_ from the front at the end of the movies (see color legends). Over time, all cells falls behind the front on average, except those that remain directly at the front until the end (for these cells, Δ_f_ > −1). To understand the dynamics of cells falling behind the front, we assumed that exterior parameters did not change over the course of the experiment and that therefore, only time differences should matter. This would imply that at any given time, the distance from the front should determine future dynamics (except for cells directly at the front). In (c) and (d), we show the distance Δ from the front of each cells (color scheme as in (a) and (b), respectively), shifted such that the final distances Δ_f_ from the front of each cell’s trajectory overlapped with the cell trajectory with the largest Δ_f_ (shown in red). Binning over intervals of 20 minutes reveals the average dynamics of cells falling behind the front (black dots): the distance Δ to the front increases exponentially (fit, dashed line) in time, independently of position, except for cell that “surf”, i.e., stay at the front for the full duration of the experiment, shown in magenta. From the shifted cell trajectories, we extracted the histograms of initial distance to the front of cells, conditional on surfing. For the histograms in Fig. 3, we pooled cell trajectories with *t* + *t*_*i*_ < 10min and *t* + *t*_*i*_ < 75min for *S. cerevisiae* and *E. coli*.

## Appendix C: Experimental methods

**1 Strains**

**S. cerevisiae – competition experiments**

To perform population growth experiments, we used strains yJHK111 (’wild type yellow strain’, [1]), yJHK112 (’wild type red strain’, [1]), and yMM9 (’red mutant strain’, unpublished, courtesy of Melanie J. I. Müller). All three strains have a W303 background (common genotype MATa *bud4*Δ::*BUD4(S288C) can1-100*, see Table C2 for details). yJHK111 expresses the yellow fluorescent protein ymCitrine, yJHK112 expresses the red fluorescent protein ymCherry. yMM9 expresses ymCherry, but is also resistant to cycloheximide (CHX) via mutation *Q37E* in gene *CYH2* (while yJHK111 and yJHK112 are sensitive). Experiments with tunable selection were performed using the pair yJHK111 and yMM9 with a variable concentration of cycloheximide in the medium. Experiments with neutral standing variation were performed using the pair yJHK111 and yJHK112. Note that, throughout this work, signal in the channel for the fluorescent color of the “mutant” (yMM9 and yJHK112) is pseudo-colored as yellow, while the fluorescent signal of the wild type (yJHK111) is pseudo-colored as blue.

**S. cerevisiae – cell tracking at front**

To track cells at the front, we used strain yMG10c, a convertant of yMG10, which in turn is based on strain yDM117 (W303 background, *HO*::*cre-EBD*, courtesy of Jasper Rine), transformed with a cassette (pMG4) based on pMEW90 (courtesy of Mary Wahl [2]). pMG4 contains a *loxP* cassette followed by ymCitrine, linked with *cyh2r* via ubiquitin. yMG10 was incubated with estradiol to induce auto-recombination, and streaked onto plates containing selective amounts of cycloheximide to select for the convertant yMG10c used for the time lapse movie in Fig. 3, which has genotype W303 *HO::cre-EBD SUC2::loxP-ymCitrine-ubq-cyh2r*.

**S. pombe**

To investigate genetic demixing from neutral standing variation in S. pombe, we used two variants of strain MJ95 (genotype *leu1*-, *ura4*-, *h*-) [3], which were obtained by replacing *mCherry* with the coding region for YFP and CFP from plasmids pOH1 and pOH2 at the *atb*2 locus.

**Plasmids with fluorescent markers cyan and yellow**

pOH1 and pOH2 are based on the vector pTrc99A, with sequences for eCFP and Venus YFP inserted between the *SacI* and *XbaI* sites, respectively. These plasmid are inducible by IPTG but we found the base level of expression of the fluorescent proteins to be sufficient without inducer. For a more detailed description see Ref. [4].

**E. coli – competition experiments**

Population growth experiments were performed using three different backgrounds:

1. DH5α transformed with pOH1 and pOH2, resulting in eOH1 and eOH2. These strains are identical to those used in Ref. [4]. For the competition experiments, we transformed eOH2 with the plasmid pA-CYC184 (New England Biolabs), conferring resistance to tetracycline, resulting in eOH3. Experiments with tunable selection were performed using the pair eOH1 and eOH3 (Fig. 4), adding low concentrations of tetracycline to the growth medium (in addition to ampicillin for plasmid maintenance). Experiments with neutral standing variation were performed using the pair eOH1 and eOH2 (Figs. 4 & 5).
2. Strain MG1655 (not fluorescent) and its derivative SJ102 (genotype MG 1655 *intC::λpR-YFP-Cmr*, courtesy of Ivan Matic), which constitutively expresses YFP and is resistant to chloramphenicol, allowing us to perform experiments with tunable selection (Figs. 4 & 5) by adding low concentrations of chloramphenicol to the growth medium. SJ102 was also used to study the dynamics of *E. coli* cells at the front (Fig. 3, SI movie 2).
3. A pair of JE 5713 [5] (cross between B6 [6] and KL228 [7, 8]), transformed with plasmids pOH1 and pOH2, giving rise to eOH4 and eOH5, were used for competition experiments with neutral standing variation (Fig. 5). These strains have been reported as *rodA* mutants but also carry a point mutation in the gene *mrdA* (Waldemar Vollmer, private communication), causing a round cell shape.

**Table C1:** Media and growth conditions used in this study. For plates, 2% w/v bacto agar was added to the media before autoclaving. Antibiotics were added after autoclaving to cooled media.

| Name | Composition | Used for... | Temperature [°C] |
| --- | --- | --- | --- |
| YPD | 20g/L peptone, 10g/L yeast extract, 20g/L dextrose | *S. cerevisiae* | 30 |
| YES | 5g/L yeast extract, 30g/L glucose, 225mg/L adenine, histidine, leucine, uracil, lysine hydrochloride | *S. pombe* | 30 |
| LB | 10g/L tryptone, 5g/L yeast extract, 10g/L NaCl | *E. coli* | 21, 37 |
| M9 | 200mL/L 5× M9 salts [9], 5g/L dextrose, 2mM MgSO<sub>4</sub>, 0.1 mM CaCl<sub>2</sub> | *E. coli* (Fig. 5) | 37 |

**Table C2:** Strains used in this study.

| Strain | Description | Genotype | Fluor. | Resistance | Reference | Fig(s). |
| --- | --- | --- | --- | --- | --- | --- |
| ***S. cerevisiae*** |  |  |  |  |  |  |
| yJHK111 | YFP WT | W303 *MATa bud4Δ::BUD4(S288C) can1-100 his3::prACT1-ymCitrine-tADH1:His3MX6* | yellow | - | [1] | 1,4,5 |
| yJHK112 | RFP WT | W303 *MATa bud4Δ::BUD4(S288C) can1-100 his3::prACT1-ymCherry-tADH1:His3MX6* | red | - | [1] | 1,4,5 |
| yMM9 | RFP MT | W303 *MATa bud4Δ::BUD4(S288C) can1-100 his3::prACT1-ymCherry-tADH1:His3MX6 CYH2::cyh2-Q37E* | red | CHX | unpublished | 1,4,5 |
| yMG10 | converter (RFP) | W303 *HO::cre-EBD SUC2::loxP-RFP-loxP-YFP-ubq-cyh2r* | red | G418, Nat | This study | - |
| yMG10c | converted (YFP) | W303 *HO::cre-EBD SUC2::loxP-YFP-ubq-cyh2r* | yellow | CHX, Nat | This study | 3 |
| ***S. pombe*** |  |  |  |  |  |  |
| MJ95 | RFP WT | *mCherry-atb2::hph, leu1-, ura4-, h-* | red | Hyg | [3] | - |
| MJ95-CFP | CFP WT | *CFP-atb2::hph, leu1-, ura4-, h-* | cyan | Hyg | This study | 5 |
| MJ95-YFP | YFP WT | *YFP-atb2::hph, leu1-, ura4-, h-* | yellow | Hyg | This study | 5 |
| ***E. coli*** |  |  |  |  |  |  |
| eOH1 | CFP WT | NEB DH5α *eCFP* (pTrc99A) | cyan | Amp | [4] | 4, 5 |
| eOH2 | YFP WT | NEB DH5α *YFP* (pTrc99A) | yellow | Amp | [4] | 4, 5 |
| eOH3 | YFP MT | NEB DH5α *YFP* (pTrc99A) + pACYC184 | yellow | Amp, Tet | This study | 4, 5 |
| MG1655 | WT | - | - | - | Laboratory stock | 4, 5 |
| SJ102 | YFP MT | MG 1655 intC::λ*P<sub>R</sub>*-*YFP-Cm<sup>R</sup>* | yellow | Cm | unpublished | 3, 4, 5 |
| JE5713 | Round cell shape | *mrdA*-mut | - | - | [5] | - |
| eOH4 | CFP round cells | JE5713 *eCFP* (pTrc99A) | cyan | Amp | This study | 5 |
| eOH5 | YFP round cells | JE5713 *YFP* (pTrc99A) | yellow | Amp | This study | 5 |

**2 Determining fitness differences as function of drug concentration**

**Liquid culture (*S. cerevisiae* only)**

Single colonies were picked and grown overnight, then diluted 1:10 in fresh media, and grown for another 3 generations to ensure growth in log phase. The resulting cultures were mixed at ratio *P*_i_ = 0.5 (measured by OD) and about 10000 cells inoculated into the wells of a 96 well plates with fresh YPD containing a range of antibiotics concentrations (3 replicates from the same initial culture per concentration). The plates were sealed and grown at 30°C overnight, then shaken vigorously for at least 1 minute. 10μl of the culture were diluted into PBS for analysis in the flow cytometer (Beckman-Coulter Fortessa X2). Every day, about 10000 cells were re-inoculated into fresh YPD to passage the cells for a total of 5 days, corresponding to about 60 generations. The cultures diluted in PBS were stored at 4°C until they were analyzed using the flow cytometer at a rate of at most 10000 events per second. The resulting ratio of mutants to wild type increased exponentially with the number of generations elapsed, whence the fitness difference could be calculated from the slope of the curve in a semi-logarithmic plot.

**On plates**

Fitness differences were measured in separate experiments using the colliding colony assay, described briefly in the following, see also Ref. [1]. Two 1μl droplets, each containing one of the two strains in log phase, are placed on agar plates about 5mm apart and incubated for at least 72 hours, until a sizable interface between the resulting colonies is formed. A circle of radius *R* is manually fitted to the collision interface using the Zeiss ZEN software and the distance *l* between the inoculation centers is measured. The selection coefficient s can be calculated via

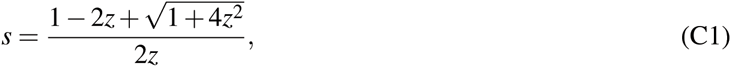
where z = *R*/*l*. The resulting values of s were found to exhibit an approximately linear dependence on drug concentration (Fig. B1a). We used the values of s given by the linear regression in the figures in the main text (Fig. 1h, i, j, Fig. 4g, h). Following the results from Ref. [1], we assumed the same regression for fitness differences on plates and in liquid culture.

**3 Adaptation from standing variation during two types of population expansions**

**Experiment and quantification**

The experimental procedure for competition experiments from standing variation is described briefly in the main text and methods therein, see also Fig. 1a for a cartoon of the experiments. Here, we provide additional details on the experimental procedures.

All competition experiments were performed on one batch of media/plates (per experimental series) and using the same overnight cultures. For each competition experiment on plates we also carried out fitness measurements (via colliding colonies) on the same batch of plates. Final population sizes of the budding yeast colonies were measured by resuspending colonies into PBS, diluting and replating to count colony forming units, or measuring optical density and comparing with a previously obtained calibration.

To measure the frequency of yeast mutants in well-mixed liquid culture, we grew and mixed the strains as described above and grew the mixture overnight in aerated culture tubes at 30°C (2 replicates from the same initial culture per concentration). We separately checked that the competed strains had the same carrying capacity to avoid error due to the cultures entering stationary phase. The next morning, we sampled 20μl from each tube into 180μl of PBS. The mixture was then measured in a Beckman-Coulter Fortessa X20 flow cytometer at a rate of at most 10000 events per second.

**Table C3:** Number of sectors analyzed per cycloheximide concentration.

|  |  |  |  |  |  |  |
| --- | --- | --- | --- | --- | --- | --- |
| [CHX] | 0 | 40 | 80 | 120 | 160 | 200 |
| Sectors | 46 | 101 | 178 | 102 | 69 | 35 |

To determine the number of cells in the outer rim of the inoculum, *N*_mut_, in Fig. 4h, we measured the radius *r*_0_ of the evaporated droplet (5 colonies from the same initial culture), which was easily visible under brightfield illumination. *N*_mut_ was then calculated as *N*_mut_ = 2π*r*_0_*P*_i_. The variable *N*_sec_/*N*_mut_ in Fig. 4g hence corresponds to the probability of surfing of an individual mutant cells in the very first cell layer, assuming that the droplet rim is perfectly flat. *N*_sec_/*N*_mut_ differs from the true surfing probability (of an individual cell in the front) by a numerical factor of order 1 taking into account the irregularities of the droplet perimeter.

**Image analysis**

To measure the frequency of mutants in colonies, images of the colonies were taken with a Zeiss AxioZoom v16 fluorescence microscope at 3.5x zoom and analyzed using custom routines written in Mathematica (Wolfram Research, Inc., Mathematica, Version 10.1, Champaign, IL (2015)). Because the colonies’ fluorescence typically becomes weaker near the colony boundary, we employed a local adaptive binarization scheme. Since individual images varied in intensity distribution, it was necessary to set the binarization thresholds by hand for each image such that the binarized shape corresponded well to the observed sector shapes. We expect the error from this “subjective” choice of thresholds to be small. During binarization, the outer radius of the colony, its center, and the radius of the inner ring, stemming from the inoculation droplet, were also measured. The frequency of mutants was then calculated by measuring the area of mutants and dividing by the area of the annulus between the outer and inner radius, i.e., the fraction inside the homeland was neglected, but the emerging bulge (for larger *s*) was taken into account.

For the frequency per sector in Figs. 1 and B7, we selected only colonies that either only had a single sector, or colonies with few sectors that did not touch. Since the colonies used for Fig. 1 had many sectors at large s, we also used colonies from experiments with smaller *P*_i_ (0.0025, 0.005, 0.01) to acquire enough “free-standing” sectors. The frequency was then computed as described above. Table C3 gives the number of sectors analyzed for each concentration of cycloheximide.

**4 Growth of *S. cerevisiae* colony from single cell**

Using a Zeiss AxioZoom v16 upright microscope, we tracked the growth of a colony (strain yMG10c) by taking time-lapse movies of the fluorescence signal detected in a stage-mounted Okolab UNO-PLUS incubator at 30°C and at constant relative humidity. An agar plate in a Petri dish was inoculated with single cells and grown in the stage-top incubator until colonies were visible at the desired magnification. Then, one colony was randomly chosen and the time lapse movie was recorded for 48 hours, taking an image every 30 min. The colony radius was determined by fitting a circle to the circumference of the colony.

**5 Cell tracking at the front**

**Experiment**

For single-cell resolution time lapse movies (SI movies 1 and 2) of growing SJ102 and yMG10c, we used a Zeiss LSM700 in confocal mode with a 488nm laser. Agar plates were inoculated with fresh culture droplets (2μl), that were left to dry for several minutes. The agar around the droplet was then cut into a 2cm×2cm pad and inverted onto a clean coverslip, such that the cells touched the coverslip. The coverslip with the cells was then incubated for a day to reach steady-state growth of the colony. After mounting the coverslip in a stage-top incubator, we mounted the incubator on the microscope and let it equilibrate for about 2h. Humidity in the chamber was controlled by the addition of a water reservoir. *E. coli* cells were imaged with a 40x oil objective, *S. cerevisiae* cells with a 20x air objective. Images were taken at 1 frame per minute with a dwelling time of about 6μs/px (31s exposure per frame) for 274/228 minutes, respectively.

**Analysis**

For cell tracking, all frames were cleaned automatically using a median filter and contrast-adjusted. In SI movie 2, some frames were manually retouched to remove brightness fluctuations. All frames were segmented with a local adaptive binarization algorithm (same parameters for all frames) and objects touching the image boundaries were removed. Because cells far behind the front could usually not be tracked accurately, we only analyzed the first few cells layers by automatically finding the position of the front and removing segmented objects far from it.

To determine the ancestry of cells at the front, we proceeded backwards in time. An individual cell was tracked by creating a mask from its outline, dilating it, and computing the overlap with the previous frame. The cell’s position in the previous frame was then determined by finding the cell with maximal overlap.

For Fig. 3, we tracked a total of 692 and 407 cells for 180 minutes in *E. coli* and *S. cerevisiae*, respectively, i.e., we shortened the original time lapse movies to 180 minutes. This was done to maximize the number of tracked cells while still maintaining information over sufficiently long time scales.

To obtain the mean square displacement in Fig. 3f, we proceeded as follows. Each tracked cell in the final frame was traced back to its ancestor 180 minutes ago. Since the front had a defined direction of motion (which we defined as the x-axis), we measured, in each time step, the position of the cell relative to its original position and computed the displacement y from the *x*-axis, and take the square. These operations are performed for all tracked cells, and averaging is performed over bins of *x*-displacements to account for cells moving by different amounts in the *x*-direction per frame. After averaging, the square root was taken in each *x*-bin, and the curves were fitted using *Mathematica*. In order to compare values for *S. cerevisiae* and *E. coli*, we divided the displacements in *x* and *y* by the effective cell sizes *d*, given by *d* = 4.5μm for *S.cerevisiae* and 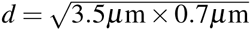 for *E.coli*. The effective cell size for E. coli was determined by the harmonic mean of its semi-axes, which were both measured directly from the time-lapse movie, as was the cell size of *S. cerevisiae*.

**Figures**

Figures of the cell tracking (Figs. 3a, c and Fig. B8) were created using Adobe Photoshop by overlaying images of the segmented cells at *t* = 0 and *t* = 3h with the computed lineages. For Figs. 3a & c, an outline was added to the tracked lineages and the cells in the lineage to increase visibility.

